# Characterization of the immunoglobulin lambda chain locus from diverse populations reveals extensive genetic variation

**DOI:** 10.1101/2022.07.20.500849

**Authors:** William S. Gibson, Oscar L. Rodriguez, Kaitlyn Shields, Catherine A. Silver, Abdullah Dorgham, Matthew Emery, Gintaras Deikus, Robert Sebra, Evan E. Eichler, Ali Bashir, Melissa L. Smith, Corey T. Watson

**Affiliations:** Department of Biochemistry and Molecular Genetics, University of Louisville, Louisville, KY; Department of Genome Sciences, University of Washington School of Medicine, Seattle, WA; Howard Hughes Medical Institute, University of Washington, Seattle, WA; Google Accelerated Science Team, Google Inc, Mountain View, CA; Icahn Institute of Genomics Technology and Data Science, Icahn School of Medicine at Mount Sinai, New York, NY; Sema4, Stamford, CT

## Abstract

Immunoglobulins (IGs), crucial components of the adaptive immune system, are encoded by three genomic loci. However, the complexity of the IG loci severely limits the effective use of short read sequencing, limiting our knowledge of population diversity in these loci. We leveraged existing long read whole-genome sequencing (WGS) data, fosmid technology, and IG targeted single-molecule, real-time (SMRT) long-read sequencing (IG-Cap) to create haplotype-resolved assemblies of the IG Lambda (IGL) locus from 6 ethnically diverse individuals. In addition, we generated 10 diploid assemblies of IGL from a diverse cohort of individuals utilizing IG-cap. From these 16 individuals, we identified significant allelic diversity, including 37 novel IGLV alleles. In addition, we observed highly elevated single nucleotide variation (SNV) in IGLV genes relative to IGL intergenic and genomic background SNV density. By comparing SNV calls between our high quality assemblies and existing short read datasets from the same individuals, we show a high propensity for false-positives in the short read datasets. Finally, for the first time, we nucleotide-resolved common 5-10 Kb duplications in the IGLC region that contain functional IGLJ and IGLC genes. Together these data represent a significant advancement in our understanding of genetic variation and population diversity in the IGL locus.

## Introduction

Immunoglobulins (IGs) or antibodies are essential protein components of the immune system that recognize and uniquely bind antigen resulting in activation and/or modulation of innate and adaptive immunity. IGs are produced by B cells and are either excreted as antibodies or expressed on the cell surface as B cell receptors (BCRs). IGs are composed of two identical heavy chains (IGH) and two identical light kappa (IGK) or lambda (IGL) chains. The role of the light chain is primarily for antigen interaction, structural stability, and autoantibody temperance(1–3). The light chain is partitioned into two primary structures: the constant domain and the variable domain. The variable domain can interact with antigen through complementarity determining regions (CDRs), while also providing structural support to the heavy chain. The light chain constant domain forms a cysteine bond with the heavy chain while providing additional structural support. The light chain variable domain is encoded by an extensive assortment of variable (V) and joining (J) genes, while the constant region is encoded by constant (C) genes. The genes encoding the V, J and C genes for IGK and IGL reside in two independent loci in the human genome, chromosome 2 (2p11.2; IGK) and chromosome 22 (22q11.2; IGL), respectively. To create a complete light chain, B cells undergo a unique process referred to as V(D)J recombination(4), during which a single V and J gene are selected from one of either the IGK or IGL loci and spliced together at the DNA level to form a complete variable region. This region is then joined with a constant (C) gene during RNA splicing to create a complete light chain. V(D)J of the light chain occurs iteratively following the production of a complete heavy chain. First, the IGK region undergoes V(D)J recombination and productive light chains are paired with the heavy chain until a suitable pairing is established. If there are no productive IGK V(D)J rearrangements, the IGL region then undergoes V(D)J recombination to produce a light chain. The ratio of IGK/IGL for naive repertoires is approximately 1.5-2; however, this ratio can differ significantly depending on the heavy chain(5).

The human IGL region spans ~920 KB of chromosome 22 (GRCh38; chr22:22,024,092-22,944,092). This region has been sequenced to completion and fully annotated twice, the first by using cosmid and BAC libraries from two different individuals by Kawasaki, et. al., in 1999. Then Watson, et al., in 2015 assembled a second haplotype using bacterial artificial chromosomes (BACs) from the hydatidiform mole cell line CHM1, and now is included as the reference IGL region in the current GRCh38 assembly(6). The GRCh38 IGL locus contains 73 variable (IGLV) genes, including 33 functional and six open reading frame (ORF) genes that are grouped into eleven phylogenetic subfamilies, as defined by the Immunogenetics Information System (IMGT). An additional functional IGLV gene, IGLV5-39, has also been described as part of a common 9.2 KB structural variant (SV)(7), which is present in the Telomere-to-Telomere (T2T) CHM13 assembly(62). There are currently 98 functional IGLV alleles curated in the IMGT database(9); however, it is clear from recent sequencing efforts that additional allelic variants remain to be described in the human population. For example, a recent study of 100 Norwegians discovered 6 novel IGLV alleles not documented in the IMGT database(10,11).

Structural diversity has also been observed outside of the IGLV gene cluster. The IGL locus constant region (IGLC) contains alternating J and C genes arranged in 3.5-5.6 KB cassettes, with each cassette containing a single J and C gene. Following V(D)J recombination, a V-J gene transcript is spliced with the C gene directly distal of the J gene(12). In the GRCh38 reference there are seven IGLJ-C gene cassettes, four of which produce functional light chains. Evidence of SVs in the IGLC region have been reported, with some individuals having 5-10 KB insertions that contain one or two additional functional IGLJ-C gene cassettes(13,14). These IGLJ-C gene cassette duplications are not present in either of the fully sequenced IGL haplotypes and the full sequence of a duplication haplotype has not yet been described with nucleotide resolution. Akin to IGLV, allelic variants have also been described for IGLJ and IGLC genes; however, it is likely that there are additional variants to be discovered.

While there is a plethora of human whole genome sequencing (WGS) short-read data available (15), short-read approaches perform poorly in clearly resolving other IG loci(16). The high density of segmental duplications and repetitive elements in the IGL locus is comparable to other IG loci(6). These features likely limit the accuracy of standard short read sequencing approaches in detecting genomic and structural variation within the locus. More recently, adaptive immune receptor repertoire sequencing (AIRR-seq) has been used to infer IG germline variation by sequencing IG receptor transcripts(11,17–22). However, AIRR-seq based approaches are limited to characterizing germline variation in exonic and upstream untranslated regions; thus, fully resolving intergenic variants, including single nucleotide variants (SNVs) in intronic and recombination signal sequences, as well as SVs, is not possible. The limited number of IGL references and lack of adequate high-throughput methodologies to accurately resolve the IGL locus at the population scale has thus far stunted our ability to fully identify and curate germline variation in IGL. These limitations have significant downstream impact on the accurate analysis of immune repertoire datasets and disease association studies(23).

To begin to build a more comprehensive catalog of IGL haplotypes, we generated and characterized reference-quality haplotype assemblies for the IGL locus from six individuals, samples from which were originally collected as part of the 1000 Genomes Project (1KGP) cohort, and which have been profiled previously using multiple orthogonal methodologies. To augment this initial set of haplotypes, we adapted and employed a targeted single-molecule, real-time (SMRT) long-read sequencing IG enrichment method(16) to conduct high-throughput IGL haplotype characterization in an additional sample set of 10 donors from ethnically diverse populations. From this collection of haplotype assemblies, we have annotated alleles, SVs, and SNVs in these individuals, revealing a high degree of allelic and nucleotide diversity. We demonstrate elevated SNV density in IGL, particularly in gene segments. Importantly, we show that many SNVs and SVs are not effectively captured by standard short-read next-generation sequencing (NGS). Finally, our extensive characterization of these IGL haplotypes has identified multiple novel IGLV and IGLC alleles, as well as the first descriptions of IGLJ-C duplication haplotypes with nucleotide resolution. Together this study includes 32 vetted and fully-characterized IGL haplotypes, including 12 from six fully phased individuals, and seven gapless haplotypes.

## Results

### Assembly of the IG lambda chain locus

We first generated high-quality assemblies from six 1KGP individuals(24)(Fig. Methods Overview) (NA18956, NA19240, NA18555, NA12878, NA19129, NA12156) using fosmid clones and long-read sequencing data derived from selectively sequencing DNA enriched for IGL as described in the Materials and Methods. An average of 80 fosmid clones per individual (Table 1) whose end-sequences mapped to the IGL locus were sequenced using SMRT long-read sequencing on the Pacific Biosciences RSII and/or Sequel lie systems. Each fosmid was uniquely assembled and then used to construct haplotype-resolved contigs spanning the IGL locus. These initial assemblies were further augmented using contigs assembled from targeted long-read sequencing to fill any remaining gaps from the fosmid-only assemblies. Finally, these targeted HiFi reads were also used to polish the assemblies. In the cases of NA19240 and NA12878, publicly available whole genome long-read sequencing data were also used to resolve any residual gaps. Using familial data, haplotype phasing accuracy was confirmed for NA19240, NA12878, NA19128, and NA12156.

**Table 1.** Table describing the base-pair contribution of fosmid, capture, and wholegenome sequence to the final hybrid assembly.

| Sample | Library | Population | Concordant<br>Fosmids<br>sequenced | Discordant<br>Fosmids<br>sequenced | Bases added<br>using fosmids<br>(bp) | Bases added<br>using WGS (bp) | Bases added<br>using capture<br>(bp) | Number of gaps | Size of gaps<br>(KB) | Gap as percent<br>of total<br>assembly | Total assembly<br>sequence | Fosmid as<br>percent of total<br>assembly |
| --- | --- | --- | --- | --- | --- | --- | --- | --- | --- | --- | --- | --- |
| NA18956 | ABC9 | Japanese | 52 | 0 | 1279669 | 0 | 576878 | 0 | 0 | 0.00% | 1856547 | 68.90% |
| NA19240 | ABC10 | Yoruban | 94 | 7 | 1626664 | 238081 | 0 | 0 | 0 | 0.00% | 1864745 | 87.20% |
| NA18555 | ABC11 | Chinese | 94 | 5 | 1562873 | 0 | 299058 | 0 | 0 | 0.00% | 1861931 | 83.90% |
| NA12878 | ABC12 | European | 68 | 2 | 981527 | 42228 | 0 | 7 | 763.5 | 74.58% | 1023755 | 95.90% |
| NA19129 | ABC13 | Yoruban | 91 | 4 | 1599736 | 0 | 226944 | 3 | 10.6 | 0.58% | 1826680 | 87.60% |
| NA12156 | ABC14 | European | 82 | 6 | 1579363 | 0 | 62013 | 5 | 183 | 11.15% | 1641376 | 96.20% |

Among the haplotypes assembled from 6 donors, we observed a range in locus coverage (20-100%; Table S1). The majority of identified gaps were the result of V(D)J recombination, as these samples are all derived from EBV-transformed B cell lymphoblastoid cell lines (Table S1). Seven haplotypes, including those from NA18956, NA19240, NA18555, NA12156, spanned the entire IGL locus with zero gaps. Locus coverage for the remaining assemblies ranged from 99.53% in NA19129 to 20.2% in NA12878 (overall mean=92.64%). The most significant example of this was for NA12878, in which a single V(D)J recombination resulted in the loss of sequence spanning IGLV3-6 to IGLC2 for the first haplotype and >80% of the locus from the second haplotype.

To further investigate sequence diversity in IGL we used IG-Capture to sequence 10 additional individuals from the 1KGP cohort (Fig. Methods Overview). In these individuals, average read depth across IGL ranged from 105x to 453x coverage (mean=183x; Table S2). Assembly contigs generated by IGenotyper(16) covered 97.8% to 99.9% of the IGL locus for individuals with an overall average coverage of 99%. Due to V(D)J recombination artifacts, gaps of 108 KB and 203 KB were observed in one haplotype each of HG02061 and NA10831, respectively (Table S2). Average phasing block size over IG regions ranged from 36 KB (HG01258) to 119 KB (NA18508) (mean=72 Kb). Samples without V(D)J artifacts had 62% (HG01258) to 94% (NA18508) of IG regions covered by phased blocks (mean=81.75%; Table S3).

### Allelic Diversity in the IGLV Genes

We were able to haplotype resolve all functional and ORF alleles for five of the six hybrid assembly samples (Fig. 1A). Due to aforementioned V(D)J recombination associated artifacts, NA12878 was missing IGLV3-1 and IGLV4-3 on both haplotypes, and an additional 26 genes on one of the haplotypes (haplotype “2”; Fig. 1A). For the ten capture samples, we were able to fully resolve all genes from both haplotypes in eight samples. Again, due to artifacts associated with V(D)J recombination, HG02061 was missing nine genes on one haplotype and NA10831 was missing 14 genes on one haplotype (Fig. 1B).

**Fig 1.**
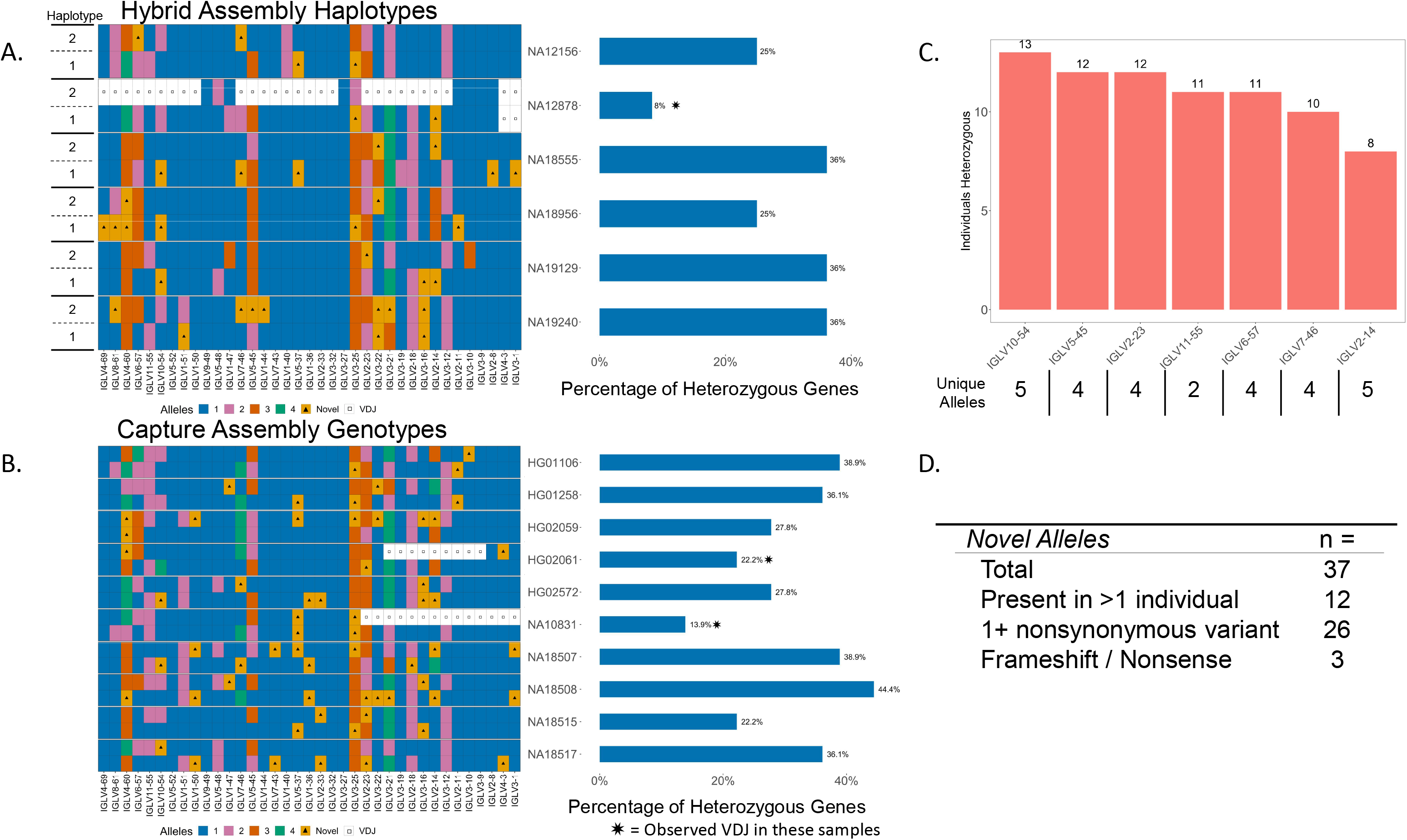
(A) Diagram of haplotype-resolved IGLV alleles from hybrid assemblies of 6 individuals (left). Colors in each tile represent allele assignments (see key), triangles denote alleles not found in the IMGT database, white tiles with a square represent unresolvable regions due to V(D)J recombination artifacts. Bar chart showing the percent of heterozygous genes in each donor (right). (B) Diagram of IGLV alleles from capture assemblies in 10 individuals with percent of heterozygous genes denoted on the right (bar chart). (C) Bar chart showing genes that are heterozygous in at least 50% of individuals; the number of heterozygous individuals observed at each gene is denoted above the bars, and the number of observed alleles at each gene is indicated (x axis labels). (D) Table of total observed novel alleles that are not present in the IMGT database, novel alleles found in >1 individual, alleles with at least one nonsynonymous amino acid substitution, and alleles resulting in frameshifted nonsense V exons.

The number of non-IMGT novel alleles identified in these assemblies ranged from three to ten per individual, with a total of 37 unique novel alleles found collectively across all samples. Of the 37 novel alleles identified, 12 were present in >1 individual (Fig. 1D). Relative to the closest IMGT allele, eight of the identified novel alleles were defined by only synonymous amino acid substitutions. Conversely, 26 novel alleles contained one or more amino acid (AA) substitutions, with 4 alleles containing 2 AA substitutions, and 2 alleles containing 3 AA substitutions. Interestingly, three identified alleles harbored insertion/frameshifts that resulted in non-productive coding sequences (Table S4, Fig. S1). IMGT annotated framework (FR) regions contained 20 of the 34 AA substitutions, and CDR regions contained 14 AA substitutions. Substitutions in the CDR regions were evenly distributed between CDR1, CDR2, and CDR3, while FR substitutions primarily occurred in FR2 and FR3, with 8 and 7 substitutions, respectively (Table S4).

We also observed high levels of heterozygosity across genes within the 16 individuals characterized. Excluding samples affected by V(D)J recombination, we found that the percent of IGLV genes at which heterozygosity was observed ranged from 22.2% to 44% (mean=33.1%) across individuals (Fig. 1A, 1B). Interestingly, seven IGLV genes were heterozygous in more than half of the individuals; specifically, the genes IGLV10-54, IGLV5-45, and IGLV2-23 were heterozygous in ≥ 12 individuals (Fig. 1C). Conversely, five IGLV genes were homozygous across all individuals (IGLV3-9, IGLV3-27, IGLV3-32, IGLV1-40, and IGLV9-49), with IGLV1-40 being notable as it is utilized at high frequencies in naive repertoires(11,25).

### Structural Variation Across the IGLV Region

The IGLV region contains a single known V gene-containing SV(7,13). This SV spans a 9.1 KB region and includes the functional gene IGLV5-39, and two pseudogenes, IGLVII-41-1 and IGLV1-41(7,13,26). Notably, this region is deleted in the GRCh38 reference. Nine of the 16 individuals in our cohort carried at least one copy of the IGLV5-39 SV (Table S5). Every haplotype with the IGLV5-39 SV contained the same allele, IGLV5-39*01. The insertion breakpoints are consistent between individuals, with every individual examined demonstrating the same breakpoint (Fig. S2). To determine if the SV was due to non-allelic homologous recombination (NAHR) we searched for homologous sequence within a 1 KB region flanking the insertion breakpoints in individuals with the insertion, but did not identify putative homologous content. The lack of homology in the sequence surrounding the SV breakpoints suggests that NAHR is not the primary mechanism of IGLV5-39 deletion. Investigating the SV breakpoints reveals a 181 bp stretch of 87.3% A/T content that is 240 bp from the proximal breakpoint (Fig. S3). As suggested by others previously, it is possible that AT-rich regions can cause replication fork slippage, which may have resulted in the IGLV5-39 deletion haplotype(27,28).

To further investigate the origin of the IGLV5-39 SV, we compared the SV sequence to the 60 KB region surrounding the insertion breakpoint and found two proximal regions and one distal region with ~6.7 KB of sequence homology surrounding the genes IGLV5-35, IGLV5-45, and IGLV5-48 (86%, 87%, and 86% sequence similarity, respectively). Additionally, there is a −14-20 KB region surrounding IGLV5-35, IGLV5-45, and IGLV5-48 that have 84-88% sequence similarity (Fig. S4). These regions surrounding IGLV5-35, IGLV5-45, and IGLV5-48 have been identified as an ancient segmental duplication event in GRCh38 by the tool SDquest(29) (Fig. S5). Using SDquest on the NA19240 haplotype containing the IGLV5-39 insertion, the insertion, surrounding region, and IGLV5-35, IGLV5-45, and IGLV5-48 were identified as ancient segmental duplications(Fig. S5). Segmental duplications are common in the IG loci and have been reported in IGL(3,6). It is likely that the IGLV5-39 SV is the result of an ancient segmental duplication involving the region surrounding IGLV5-45 and/or IGLV5-49 and was later lost, potentially due to replication fork slippage.

Additional SVs were observed in our cohort, including two separate −6 KB LINE-insertion haplotypes, the first adjacent to IGLV1-47 and the second adjacent to IGLV4-60 and IGLV8-61 (Table S6). Further, we observed multiple regions with small SVs ranging in size from 175-320 bp (Table S7, Fig. 6S A-D). While some of these SVs are adjacent to functional genes, none are predicted to interfere with functionality.

### Single Nucleotide Variation across the IGLV region and within IGLV genes

We then characterized SNVs across the assemblies by identifying nucleotide differences relative to the GRCh38 reference spanning the 920 KB IGL region (chr22:22,024,092-22,944,092). To better characterize variant calls, the IGL region was further divided into three regions, IGLV (chr22:22024092-22886736, 862,644 bp), a 594,267 bp subset of the IGLV region, deemed IGLV IG-only, which excludes non-IG gene regions (Fig. 2A), and IGLC (chr22:22886736-22944092, 57,356 bp) which was excluded from this analysis as we chose to focus specifically on the IGLV region. These non-IG gene regions are devoid of IGLV genes, and harbor the genes VPREB1, ZNF208A, ZNF208B, and GGTLC2. Three of the 16 individuals (NA12878, NA10831, HG02061) had moderate or significant V(D)J recombination associated artifacts affecting one or both haplotypes and were excluded from SNV analysis. NA18555 had the highest number of SNVs, with 3078 across the IGL locus, 1800 of which were within IGLV IG-only regions. HG01258 had the lowest SNV count with 1351 across the IGL locus and 868 in IGLV IG-only regions (Table S8). The majority of SNV positions were heterozygous in each sample, with a mean of 65.4% (54.3% for HG02059 and 80% for NA18517); remaining positions were homozygous for an alternative nucleotide (Table S8). Across the 13 samples, there were 7599 unique SNVs in IGLV with 4298 unique SNVs in IG-only regions. Of the 4298 unique IG-only SNVs, 939 (21.8%) were not found in the dbSNP “common” database, while 124 appeared to be completely novel and not found in any dbSNP database (Table S9).

**Fig 2.**
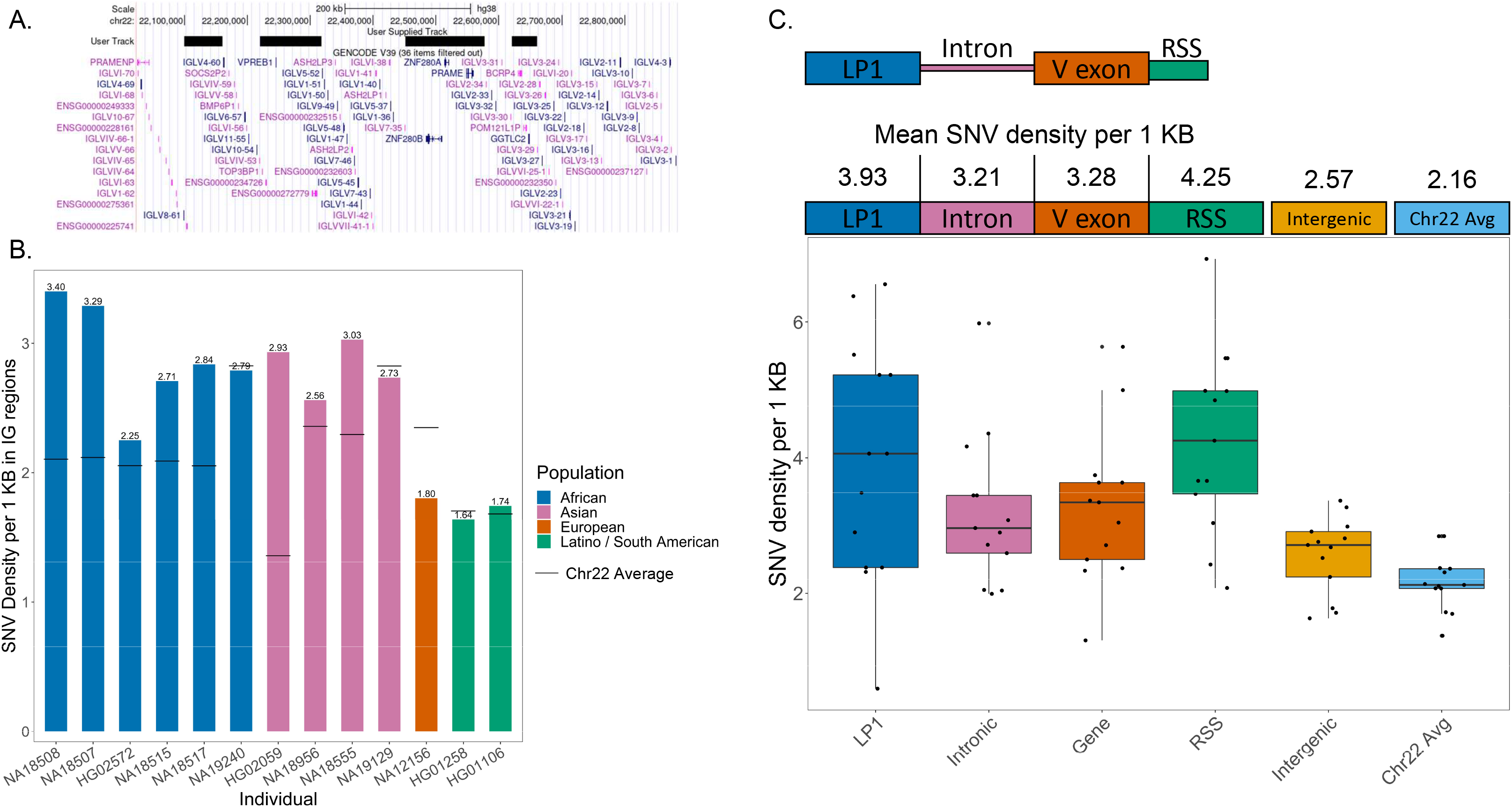
(A) Overview of the IGLV region with black bars covering coordinates masked for the “IG-only” region. (B) Density of SNVs per 1 Kb in IG-only regions are reported for 13 individuals which did not have significant VDJ artifacts. The solid black lines are average SNV density across a 22.9 Mb segment of Chr22 (q11.21-q13.31). (C) Density of SNVs in LP1, Intron, V exon, RSS, IGLV intergenic sequence, and across q11.21-q13.31 of Chr22 was measured. (D) IGLV and IGLC SNVs detected from capture and hybrid assemblies were compared to SNVs in the 1KG 30x coverage datasets. The blue bar represents SNVs called in both datasets, while green is SNVs unique to the 1KG dataset and pink is SNVs unique to the capture/hybrid assemblies. Only regions covered by capture/hybrid assemblies were considered. IGLV samples marked with stars contain the 9.2 Kb IGLV5-39 insertion while IGLC samples marked with stars contain either a 5.5 Kb or an 11 Kb duplication in the constant region. (E) A table showing SNVs only found in the 1KG 30x coverage dataset that are in V exons. Unique SNVs are the total number of non-redundant SNVs in each V exon. Individuals affected are the number of individuals with at least 1 SNV in a V exon.

We calculated the density of SNVs per 1 KB in the IGLV IG-only regions and compared this to the average background SNV density of chr22 within a 22.9 MB window (q11.21-q13.31). We chose this region specifically to represent basal chromosomal variance background, as it contained no assembly gaps in the GRCh38 reference assembly. Nine of the 13 samples had a higher density of SNVs in the IGLV IG-only region when compared to the chr 22 background (Fig. 2B). We further examined SNV density in the components of V gene segments, including leader part 1 (LP1), introns, V exons, and recombination signal sequences (RSS). Interestingly, LP1, introns, V exons, and RSS sequences on average have higher density of SNVs when compared to both the chr22 background and the IGLV intergenic space (Fig. 2C). On average, SNV density in gene components was 43% higher than IGLV intergenic sequence, and 70% higher than the chromosome 22 background.

### IGLC Haplotype Diversity

The IGL constant region of the GRCh38 reference(6) contains seven J and C gene pairs (Fig. 3A). IGLJ genes are 38 bp in length, while IGLC genes vary from 304 bp to 322 bp; with all functional C genes being 318 bp. Each J and C gene pair is part of a larger cassette, referred to here as an IGLJ-C cassette. Cassettes IGLJ-C1, IGLJ-C2, IGLJ-C3, and IGLJ-C7 are functional, whereas IGLJ-C4, IGLJ-C5, and IGLJ-C6 are nonfunctional. Both IGLJ-C4 and IGLJ-C5 contain deletions that disrupt the RSS site of their respective J genes while IGLC5 contains an 11 bp deletion, which renders it nonfunctional. IGLJ-C6 is able to participate in recombination, but the IGLC6 gene contains a premature stop codon, resulting in a truncated protein. Each cassette breakpoint is flanked by homologous sequence, a characteristic feature of NAHR. IGLJ-C1, IGLJ-C2, and IGLJ-C3 cassettes range in size from 5300 bp to 5900 bp, while IGLJ-C4 through IGLJ-C7 are 3200 bp to 3900 bp. The GRCh38 reference contains one copy of each IGLJ-C cassette (Fig. 3A).

**Fig 3.**
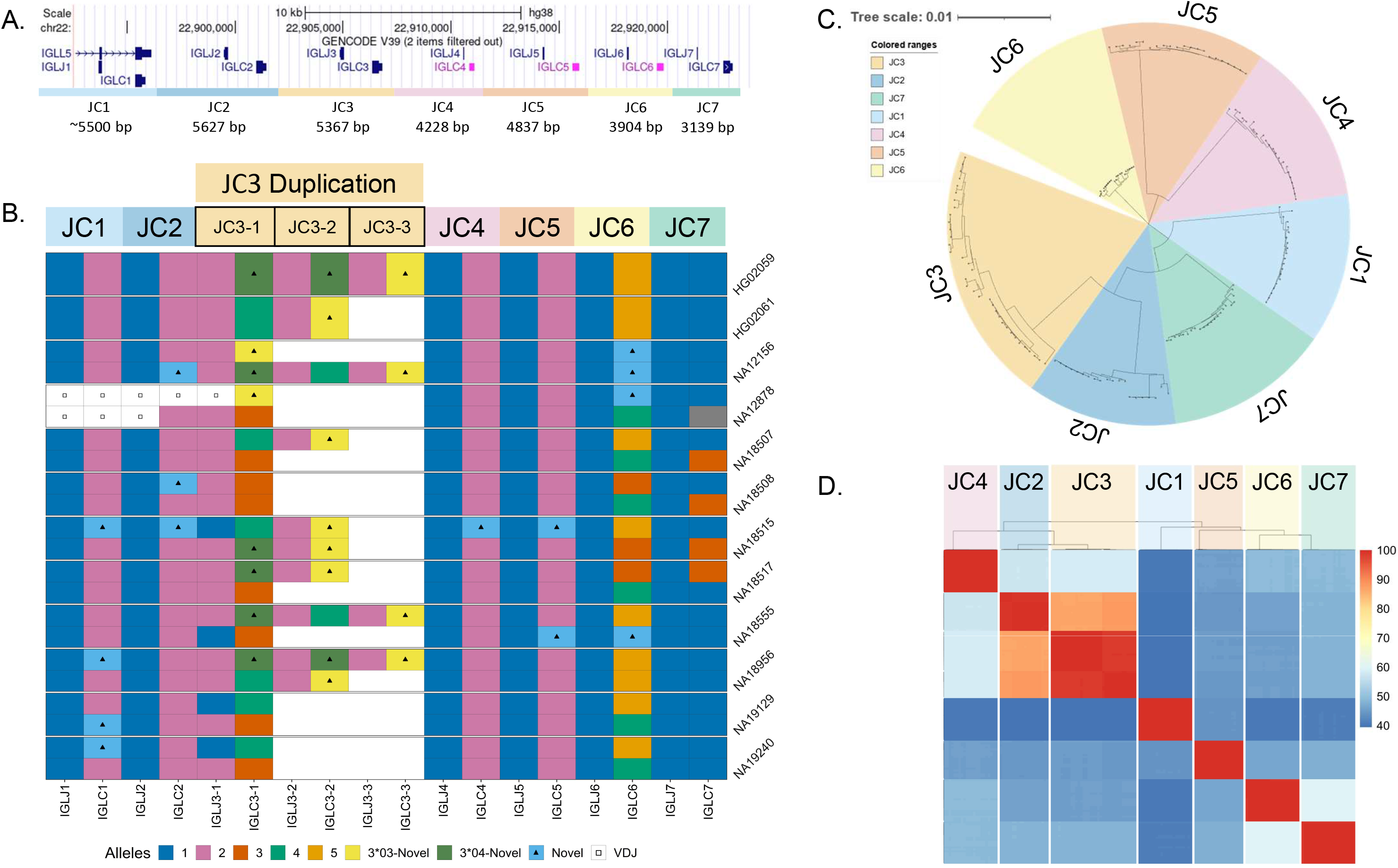
(A) A UCSC Genome Browser schematic of the IGLC region in GRCh38 (chr22:22,892,572-22,923,863). The region is divided into seven cassettes, each cassette contains a single IGLJ and IGLC gene. Cassettes JC4, JC5, and JC6 are nonfunctional. (B) Diagram of IGLC alleles for 22 resolved haplotypes. Haplotypes have either one, two, or three copies of the JC3 cassette, with blank space denoting no duplication present. In the JC3 region, two separate colors with triangles denote two separate novel IGLC alleles (3*03-Novel, 3*04-Novel). (C) Neighbor-joining tree of all resolved JC cassette sequences was generated using MSA (MUSCLE), the Neighbor-joining tree building method and the Tamura-Nei nucleotide substitution model to demonstrate sequence groups within and between JC cassettes. (D) A comparison of percent identity between all cassette sequences organized by phylogeny.

Previous studies have reported duplications of the IGLJ-C2 and IGLJ-C3 gene regions(13), with duplication cassettes having been individually Sanger sequenced using “gene walking” methodologies(14). Analysis of the IGLC haplotypes resolved from our cohort revealed significant variation in this region. Our haplotypes included one to three copies of the IGLJ-C3 cassette, which was further associated with ~5.4 KB and −10.7 KB of additional sequence (Fig. 3B). Eight of the 22 IGLC haplotypes carried a IGLJ-C3 duplication. Considering both IGLJ-C3 IGLJ/IGLC allelic variants and copy number, we characterized nine unique IGLJ-C3 haplotypes, four with a single IGLJ-C3 copy and four with multiple IGLJ-C3 copies. All IGLJ-C3 duplication haplotypes had >1 unique IGLC3 allele containing one or more nonsynonymous nucleotide variants. Notably, NA18956 carried two different duplication haplotypes, including a single copy of the IGLC3*04 allele, and two copies each of the novel alleles, IGLC3*03-Novel and IGLC3*04-Novel (Fig. 3B). Across all resolved IGLJ-C3 cassettes, we observed two IGLJ3 alleles and four IGLC3 alleles, two of which were novel. Additionally, we identified two functional novel IGLC1 alleles harboring nonsynonymous substitutions, with one allele appearing in three individuals. Two functional novel IGLC2 alleles were also identified, one of which harbored a nonsynonymous substitution and the other matching the sequence of IGLC3*04. Finally, we identified five novel pseudogene alleles, one in IGLC4, two in IGLC5, and two in IGLC6. Overall we identified 11 novel IGLC alleles, six in functional genes and five in pseudogenes; this total represents a 50% increase in total IGLC alleles known to date.

We compared the similarity of each resolved IGLJ-C cassette sequence using multiple sequence alignment and phylogenetic analyses(30)(Fig. 3C & 3D). The phylogenetic tree in Fig. 3C revealed that the IGLJ-C2 and IGLJ-C3 cassettes are most similar, sharing ~85-90% sequence identity over ~5.5 Kb. While all IGLJ-C3 duplication blocks in each haplotype are highly similar (>98%)(Fig. 3D), there were two sub-clades identified within the larger IGLJ-C3 clade (Fig. S7). The IGLJ-C3 cassettes that are adjacent to a IGLJ-C2 cassette (proximal) form a sub-clade, while the distal IGLJ-C3 cassettes bordering the IGLJ-C4 cassette form a separate sub-clade. IGLJ-C3 cassettes within the proximal clade differ from one another by only 0-6 nt (>99.89%) across 5.6 Kb. IGLJ-C3 cassettes within the distal clade differ by 0-33 nt (>99.38%). This is in contrast to differences observed between members of the two sub-clades, which differ by 62-85 nt (98.03-98.84%)(Fig. S7). Beyond sequence identity, both IGLJ-C2 and IGLJ-C3 cassettes have more nucleotide diversity than the others. When comparing sequence similarity within each cassette group (e.g., comparing between only IGLJ-C2 cassettes), IGLJ-C2 and IGLJ-C3 have the largest disparity within cassette groups at 98.722% and 98.025%, respectively. Comparatively, all other cassettes have >99% sequence identity within their respective groups (Table S10).

Further, the IGLJ-C2 and IGLJ-C3 cassette groups were observed to have high sequence homology with each other at ~85-90%. Specifically, a ~2 KB high homology region stretching from 150 bp proximal of IGLJ2/3 to 190 bp distal of IGLC2/3 are >99% similar between the two cassette groups (Fig. S8). Due to the high degree of sequence similarity, it is possible that the IGLJ-C2/3 duplications are a result of NAHR(31), which has been reported in other IG regions harboring segmental duplications(32). When we applied phylogenetic analyses to this homology region, we found that homologous segments of similar duplication haplotype and location would primarily cluster together (Fig. S9). Additionally, elevated GC content is observed in this homology region (average 57.8%) that is significantly higher than the average for Chr22 (47%) and genomic average (41.9%)(33). High GC content is often a signature of repeated gene conversions due to GC-biased repair of A:C and G:T mismatches(34). Taken together, these findings suggest that a combination of NAHR and gene conversion may maintain the high homology between IGLJ-C2 and IGLJ-C3.

### IGL SNV 1KGP comparison

Previously, Rodriguez et al., have demonstrated the accuracy of long-read assemblies in IG regions and highlighted discrepancies between SNVs called from long-read assemblies of these regions and those from short-read datasets, such as the 1KGP 30x high-coverage SNV call sets. To evaluate our data as compared to matched high-coverage SNV call sets, we compared capture and hybrid assembly IGL SNVs of the 13 individuals under examination to their respective 1KGP 30x high-coverage SNV call sets (Table 2–4)(15). Non-IG regions were masked for the IGLV comparisons and only regions which were covered by capture or hybrid assemblies were considered. Overall, 1KGP SNV calls differed greatly from capture and hybrid assembly calls. The highly accurate hybrid assemblies called between 87 (NA12156) and 145 (NA18555) unique SNVs in IG-only regions that were not present in their respective 1KGP datasets. Additionally, the 1KGP dataset falsely called between 27 (NA19129) and 433 (NA19240) SNVs in these regions. Analysis of capture-based assemblies revealed similar discrepancies with the 1KGP short-read based variant call sets. Capture assemblies had between 27 (NA18515) and 103 (HG02572) SNVs not found in the 1KGP dataset, while the 1KGP dataset called between 31 (NA18508) and 532 (HG01258) SNVs not present in the capture assemblies (Table 2). Notably, six of the eight individuals with >190 1KGP-unique SNVs carry the 9.1 KB IGLV5-39 insertion. Indeed, a large fraction of 1KGP-unique SNVs fell within and around the 14-20 KB homology regions spanning IGLV5-45, IGLV5-48, and the IGLV5-39 insertion regions annotated in Fig. S4. When we assessed the locations of false-positive SNVs in the 1KGP datasets, we found that each individual had at least one V exon harboring a false-positive SNV. Overall, 11 genes were affected across the 13 individuals. IGLV5-48 and IGLV5-45 have the highest number of erroneously called SNVs with 27 and 21, respectively; false SNVs in these two genes were observed in 10 and 7 individuals, respectively (Table 3). Additionally, IGLV5-45 and IGLV5-48 harbored 17 false-negatives, where the 1KGP datasets were unable to determine the true haplotype.

**Table 2.**
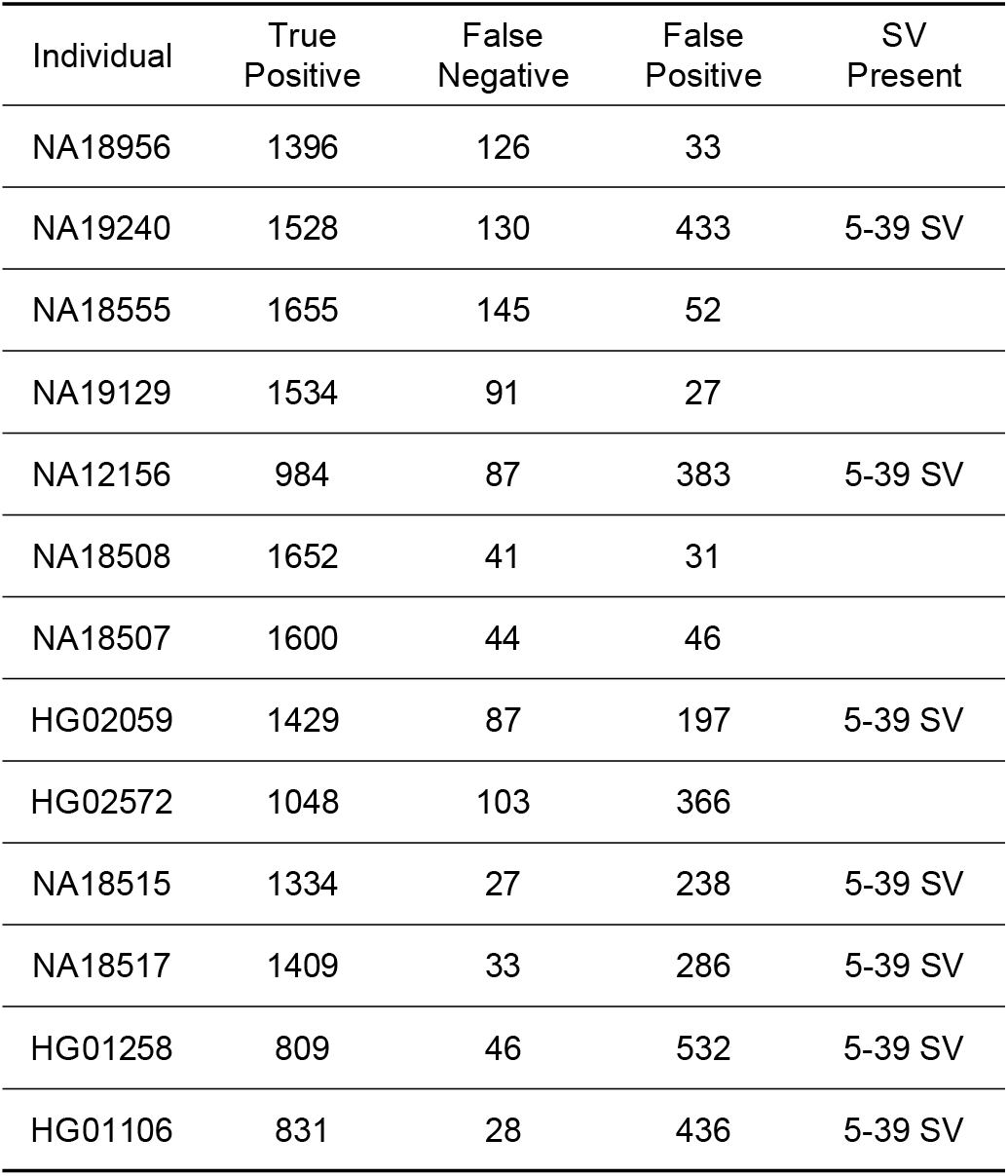
True positive, false negative, and false positive SNVs in the 1KGP dataset compared to the capture and hybrid assemblies across the IGLV region. 5-39 SV refers to the presence of a 9.1 KB insertion containing the gene IGLV5-39.

**Table 3.**
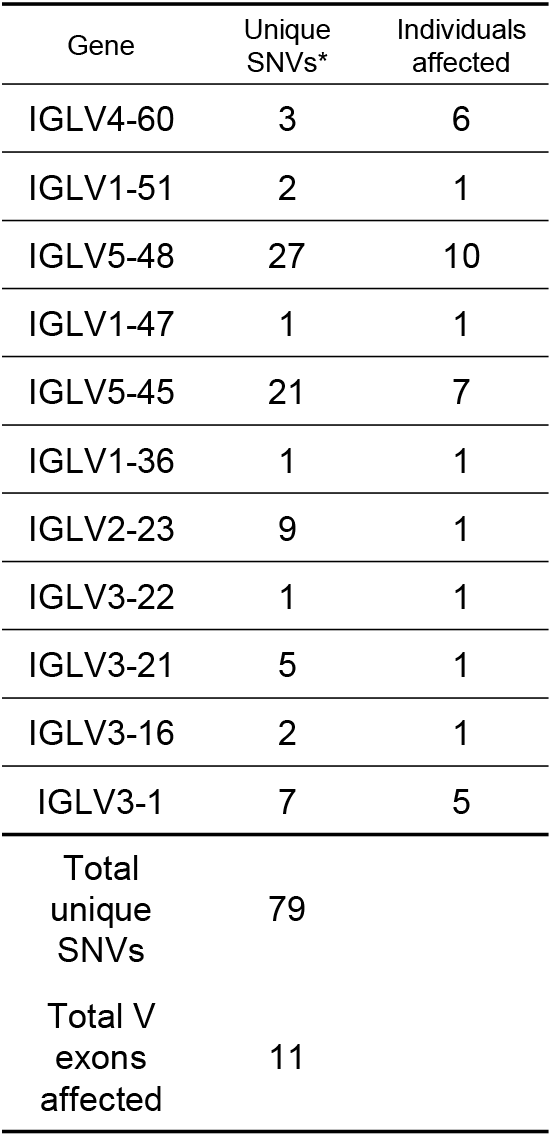
Table of genes with false positive SNVs present in the V exon as reported by the 1KGP dataset. Unique SNVs* are unique positions in each gene.

| Gene | Unique SNVs* | Individuals affected |
| --- | --- | --- |
| IGLV4-60 | 3 | 6 |
| IGLV1-51 | 2 | 1 |
| IGLV5-48 | 27 | 10 |
| IGLV1-47 | 1 | 1 |
| IGLV5-45 | 21 | 7 |
| IGLV1-36 | 1 | 1 |
| IGLV2-23 | 9 | 1 |
| IGLV3-22 | 1 | 1 |
| IGLV3-21 | 5 | 1 |
| IGLV3-16 | 2 | 1 |
| IGLV3-1 | 7 | 5 |
| Total unique SNVs | 79 |  |
| Total V exons affected | 11 |  |

For the IGLC region, we compared SNV calls from hybrid assemblies to the 1KGP datasets for the same individuals. The discrepancy between the datasets was more prominent in the IGLC region, in which the false-negative and -positive calls comprised a mean of 19.4% and 23.6% of genotypes in the 1KGP dataset (Table 4). This elevated false-positive and -negative rate is likely due to the highly repetitive and structurally variable nature of the IGLC region (Fig. 3B & 3D). In addition, NA18555, NA18956, and NA12156 contain IGLJ-C3 duplications in the IGLC region which are not present in the GRCh38 reference assembly. These results suggest that the lack of a representative reference containing multiple IGLJ-C segmental duplications results in an abundance of miscalled 1KGP SNVs.

**Table 4.** True-positive, false-negative, and false-positive SNVs in the 1KGP dataset compared to hybrid assemblies across the IGLC region. The presence and copy number of IGLJ-C3 duplications are also indicated.

| Individual | True<br>Positive | False<br>Negative | False<br>Positive | SV<br>Present |
| --- | --- | --- | --- | --- |
| NA18956 | 123 | 23 | 36 |  |
| NA19240 | 126 | 22 | 17 | J-C3 x 2 |
| NA18555 | 140 | 15 | 18 | J-C3 x 2 |
| NA12878 | 75 | 19 | 20 |  |
| NA19129 | 122 | 29 | 34 |  |
| NA12156 | 109 | 27 | 39 | J-C3 x 2 |

## DISCUSSION

Additional genomic resources and high-throughput methods for accurately resolving the IGL locus are crucial for further exploring associations between germline polymorphism, the expressed antibody repertoire, and clinical phenotypes (Watson, Glanville, Marasco 2017; Collins et al. 2020; Ivana et al.). In this study, we have characterized haplotype-resolved IGL assemblies from 6 individuals and diploid assemblies from an additional 10 individuals of diverse genetic ancestry, using a combination of long-read sequencing based datasets. These assemblies represent a significant advance, increasing the number of available vetted curated haplotype references by 32, haplotype-resolved references by 12, and 100% gapless assemblies by seven (a 3.5-fold increase). Critically, the use of long reads allowed for the comprehensive characterization of genetic diversity in these regions effectively and accurately. Moreover, we clearly demonstrate that the targeted IGLV sequencing approach we have developed and present here will allow for scalable high-throughput long-read sequencing that will enable characterization of IGL polymorphism at the population-level.

Collectively, these assemblies resolved an abundance of novel allelic variants, SNVs, indels, and SVs. Importantly, this included the identification of 37 novel IGLV alleles, six novel functional IGLC alleles, and revealed patterns of high heterozygosity and diversity. The degree of allelic heterozygosity observed here is similar to other immune loci(6), suggesting that IGL may also be under evolutionary pressures to favor elevated allelic diversity. Interestingly, there are conserved regions of homozygosity and heterozygosity across genes in the locus. For example, we found that 7/39 IGLV genes were heterozygous in >50% of individuals, while 5 others were consistently homozygous across all individuals. Patterns of high allelic diversity, elevated polymorphism, and excess nonsynonymous variation are hallmark signatures of balancing selection in the HLA genes(35). The presence of these patterns in the IGL genes, specifically among commonly heterozygous genes, suggest that these genes are also under balancing selection. In contrast, commonly homozygous genes may be under purifying selection due to specific interactions between heavy and light chains that favor preserving a specific light chain structure. Multiple forms of selection acting on a single locus has been observed before in the HLA genes, as external sources (pathogens(35,36)) and internal sources (KIR coevolution(37)) both drive evolution of this region. The lambda light chain fulfills a dual purpose role of antigen interaction and neutralization of autoreactive heavy chains(1,2). This multifunctionality may result in multiple types of selection acting on the IGL locus, balancing selection due to external antigen interactions, and purifying selection due to coevolution with IGH. As additional haplotypes are generated and curated, the application of formal tests for selection will be possible.

We also observed striking differences between coding and noncoding regions of the IGL locus. Our data show that polymorphism density is higher in IGLV gene components compared to intergenic regions, as well as the chr22 average. In addition, the majority of novel alleles contain at least one nonsynonymous substitution and overall have a higher rate of nonsynonymous to synonymous substitutions. This pattern of increased allelic diversity and polymorphism in gene coding regions has also been seen in HLA(38), further suggesting that IGLV genes are under evolutionary selection.

In addition to high levels of diversity in the IGLV region, we were able to characterize complex segmental duplications in the IGLC region. While insertions >5 KB in the IGLC region have been reported in the past(13,39) and partially sequenced(14), we present the first completely sequence-resolved IGLC duplication haplotypes. While each duplication haplotype demonstrates an exceedingly high degree of sequence similarity between duplication cassettes, the few polymorphisms that exist result in nonsynonymous differences in the constant genes. This means that individuals that carry these SVs can vary significantly in the number of functional IGLC genes. Specifically in our cohort, we see that functional IGLJ-C cassettes vary in number from four to six per haplotype. It has been observed that IGLJ-C duplication decreases IGK/IGL usage ratio, resulting in an increased utilization of lambda light chains(14). Since IGLJ-C duplication increases overall lambda light chain utilization, it is feasible that IGLJ-C duplication may introduce bias in chromosome usage during V(D)J recombination to favor the haplotype with additional gene copies. Additionally, the duplication haplotypes are also most commonly found in Asian and African super-population groups(13,14) which may suggest environmental or regional factors impacting the selection of these haplotypes. While pathogens are frequent drivers of selection(63) and gene duplications are known to be an important source of genetic novelty and adaptive evolution(40,41), the IGLC duplications have yet to be examined at population scale for signatures of selection. It will be important to further study the effect of IGLC duplications on IGHV and IGLV/IGKV pairing in the expressed repertoire.

Moving forward, it is critical that we provide a method for characterizing IGL that is accurate, scalable, and able to overcome the limitations of existing short-read sequencing approaches. Here we demonstrate the use of targeted capture and long-read sequencing to generate complete and accurate variant call sets that are able to detect and account for complex SVs, avoiding a major pitfall of short reads. This demonstrates that as additional long-read whole-genome assemblies are generated, for example, through efforts underway by the Human Pangenome Reference Consortium(61) and the T2T projects(62), these assemblies can be readily curated and integrated into IG genomic resource efforts. However, in the short term, we expect that the cost of WGS approaches will likely limit the extent of IG diversity that can be described across the human population.. Critically, our method provids a scalable and cost-effective approach for studying IGL variation at population scale at comparable resolution to WGS, but in much larger cohorts. This will be important for understanding population variation and evolution in the IG loci. Together, these data and methods advance our understanding of IG genetics and are necessary to further understand repertoire development/dynamics and antibody function in disease with genomic context.

## METHODS

### Fosmid Isolation, Sequencing, and Assembly

Fosmid clones were procured from the Human Genome Structural Variation Clone Resource (University of Washington, Department of Genomic Sciences; Seattle, WA)(24,31,43). Fosmid clones were selected based on end sequences which mapped to the IGL locus(GRCh38; chr22:22,004,092-22,964,092). For each individual, five non-overlapping pools of fosmids were selected. On average, pools contained 19 fosmids. Fosmids were grown in 96-well plates and DNA extracted following previously published methods(24). On a per fosmid clone basis, DNA was sheared using g-tubes (Covaris, Woburn, MA, USA) to −15 Kb, followed by size selection of >10 KB fragments using the BluePippin system (Sage Science, Beverly, MA, USA). Following shearing and size-selection, DNA from non-overlapping fosmids was pooled in equimolar ratios, with numbers of samples per pool dependent on the SMRT sequencing platform used. Pooled DNA was then directly prepared for sequencing using SMRTbell Template Preparation Kit 1.0 (for RSII-sequenced samples) or 3.0 (for Sequel lle-sequenced samples) (Pacific Biosciences, Menlo Park, CA, USA) following the manufacturer’s standard protocol. SMRTbells were sequenced on either the RSII sequencing system (Pacific Biosciences) using P6/C4 chemistry and 6 hour movies, or the Sequel lie system (Pacific Biosciences) using 2.0 chemistry and 30h movies.

For each fosmid pool, sequencing data from individual fosmids were used to generate fosmid-specific assemblies by first removing the fosmid backbone sequence from the reads and mapping them to the GRCh38 reference. Then, for each fosmid, only reads that mapped within the fosmid end sequence coordinates were *de novo* assembled using Canu(44).

### Capture Sequencing and Assembly

To specifically enrich for IGL we designed a custom Roche HyperCap DNA probe panel, which included target sequences from the GRCh38 IGL locus(chr22:22,004,092-22,964,092) and the GRCh38 chr22_K1270875v1_alt as targets. Genomic DNA samples were procured from Coriell Repositories (1KGP donors). Samples were prepared and sequenced following previously published methods(16). Briefly, genomic DNA was sheared to ~8 KB using g-tubes (Covaris, Woburn, MA, USA) and size-selected with the Blue Pippin system (Sage Science, Beverly, MA, USA), collecting fragments that ranged in size between 5-9 Kb. Following size selection, sheared DNA was ligated to universal adapters and amplified. Amplified, individual samples were pooled in groups of eight prior to IGL-specific enrichment using custom Roche HyperCap DNA probes following a modified protocol(16). After probe capture, targeted fragments were amplified a second time and prepared for SMRT sequencing using the SMRTbell Template Preparation Kit 3.0 following the standard manufacturer’s protocol. Resulting SMRTbell libraries were pooled in 24-plexes and sequenced using one SMRT cell 8M on the Sequel lie system using 2.0 chemistry and 30 hour movies. Following data generation, circular consensus sequencing analyses were performed, generating high fidelity (“HiFi”, > Q20 or 99.9%) reads for downstream analyses. Assemblies from capture sequencing were generated from IGL-specific reads (excluding off target reads) using IGenotyper(16) with the commands ‘phase’, ‘assemble’, and ‘detect’.

### Construction of Hybrid Assemblies

Overlapping fosmid sequences were merged by joining fosmid sequences at overlapping junctions to create continuous haplotypes. When available, parental data was used to phase fosmid haplotypes. Gaps in the fosmid haplotype assemblies were manually filled using capture assembly data from the same individual. Assemblies were then error corrected using the Inspector package(45) with HiFi reads from the capture sequencing datasets using the commands ‘inspector.py -c <assembly.fasta> -r <ccs.fastq.gz> -o output/ --datatype hifi’ followed by ‘inspector-correct.py -i output/ --datatype pacbio-hifi -o output/’. Additionally, publicly available whole genome sequencing assemblies using long-reads sequenced on the RSII sequencing system were used for error correction of NA19240(46) (GCA_001524155.4) and NA12878 (Direct Submission, Wilson,R.K. and Fulton,R.) (GCA_002077035.3).

### Identification of Novel Alleles

Alleles were extracted from completed hybrid or capture assemblies using IGenotyper(16) and annotated by comparing sequences to the IMGT/GENE-DB(9) retrieved on 2021/08/21. Novel alleles were identified through comparison to the IMGT database using IMGT/V-QUEST(47,48). Putative novel IGL alleles initially annotated from each assembly were confirmed by aligning HiFi reads to curated assemblies and assessing HiFi read support for SNVs representing each novel allele on a per sample basis; conservatively, >12 unique HiFi reads were required to confirm all novel alleles.

### Detection of Indels and SNVs from Assemblies

Variant calls were made from each assembly using IGenotyper(16). The IGenotyper ‘detect’ option calls indels and SNVs, annotating each call with a quality score, HiFi read support, and additional information such as if the indel or SNV falls in a gene or SV. IGenotyper outputs this information as a Variant Call Format (VCF) file. Using bcftools(49), the IGenotyper vcf was filtered to exclude indels and SNVs that lacked read support. The vcf file was then split using rtg-tools(50) with the ‘--snps-only’ and ‘--indels-only’ options to produce a SNV-only and an indel-only vcf file.

### Detection and Annotation of SVs from Assemblies

SVs greater than 50 bp were detected by first aligning assemblies to the GRCh38 reference, followed by visualization in the Integrated Genome Viewer (IGV)(51–53) where SVs were visually inspected. SVs greater than 1000 bp had their sequences extracted from the IGV browser using the ‘copy insert sequence’ feature. Nucleotide BLAST(54) with the Nucleotide collection (nt) database was used to identify and annotate LINE SVs. For non-LINE SVs, genes were annotated using Nucleotide BLAST for the command line(55–57) using the following parameters ‘blastn -query <assembly.fasta> -subject <gene_reference.fasta> -perc_identity 95-culling_limit 1 outfmt “6”‘. The <gene_reference.fasta> used is the IMGT/GENE-DB(9) IGL gene database retrieved on 2021/08/21. For SVs less than 1000 bp, sequence was extracted spanning ±1000 bp from the site of the SV. Extracted sequence was then aligned by multiple sequence alignment (MSA) (MUSCLE(58)) to GRCh38 at coordinates ±1000 bp from the SV. Sequence alignments were visualized using Geneious Prime (Geneious Prime 2022.11.0.(https://www.geneious.com) and annotations were added manually.

### Publicly Available SNV Datasets

SNV genotype callsets for all individuals were downloaded from the 1000 Genomes FTP portal(59). These callsets were previously generated using high coverage 30x PCR-free whole-genome shotgun short-read sequencing on an Illumina Novaseq 6000 using 2x 150 bp cycles(15). Following download, the IGL region was extracted from the SNV genotype callsets using ‘bcftools view-output-type v regions 22:22,024,092-22,944,092 -min-ac 1 -types snps’. Genotype callsets were then split using rtg-tools(50) with the ‘--snps-only’ option to produce a SNV-only vcf file.

### Calculation of SNV densities in IGLV

The background density of SNVs across Chr22 for each individual was calculated using their 1KGP high-coverage SNV data by taking the number of unique SNVs in the range chr22:19,990,736-49,936,760 and dividing by the total region size (29,946,024 bp). This region was chosen as there are no assembly gaps in the GRCh38 reference across the region. For gene components (RSS, intron, gene, LP1), the SNV density was calculated for each individual as the total number of unique SNVs across both haplotypes divided by the region’s total sequence length for each component. Only gene components of functional genes and ORFs were included in this analysis. For intergenic regions, the SNV density was calculated as the total number of SNVs located in IGLV regions (Table S12) excluding gene components divided by the region’s total sequence length.

### IGLC Cassette Clustering and Phylogenetic analysis

Sequences were first aligned by MSA (MUSCLE(58)). Geneious Prime (Geneious Prime 2022.11.0.(https://www.geneious.com)) was used to create the phylogenetic tree from the MSA using the Neighbor-joining tree building method(60) and the Tamura-Nei nucleotide substitution model(61). The phylogeny was then visualized using the online tool iTOL(62). Pairwise nucleotide similarities were estimated using Clustal Omega(30).

### Comparing 1000 Genomes Project SNVs to Assembly SNVs

SNVs detected in capture and hybrid assemblies were compared to SNV genotype callsets from the 1KGP high-coverage dataset(15). Overlap of SNVs was calculated using rtg-tools with the following command ‘rtg vcffilter --snps-only’ and ‘rtg vcfeval’. SNVs were only considered concordant when the genotype called in both datasets was the same.

## ACKNOWLEDGEMENTS

This work was supported, in part, by US National Institutes of Health (NIH) grant HG010169 to E.E.E.

E.E.E. is an investigator of the Howard Hughes Medical Institute.

## CONFLICT OF INTEREST STATEMENTS

E.E.E. is a scientific advisory board (SAB) member of Variant Bio, Inc.

Robert Sebra is VP of Technology Development at Sema4.

## Methods Overview

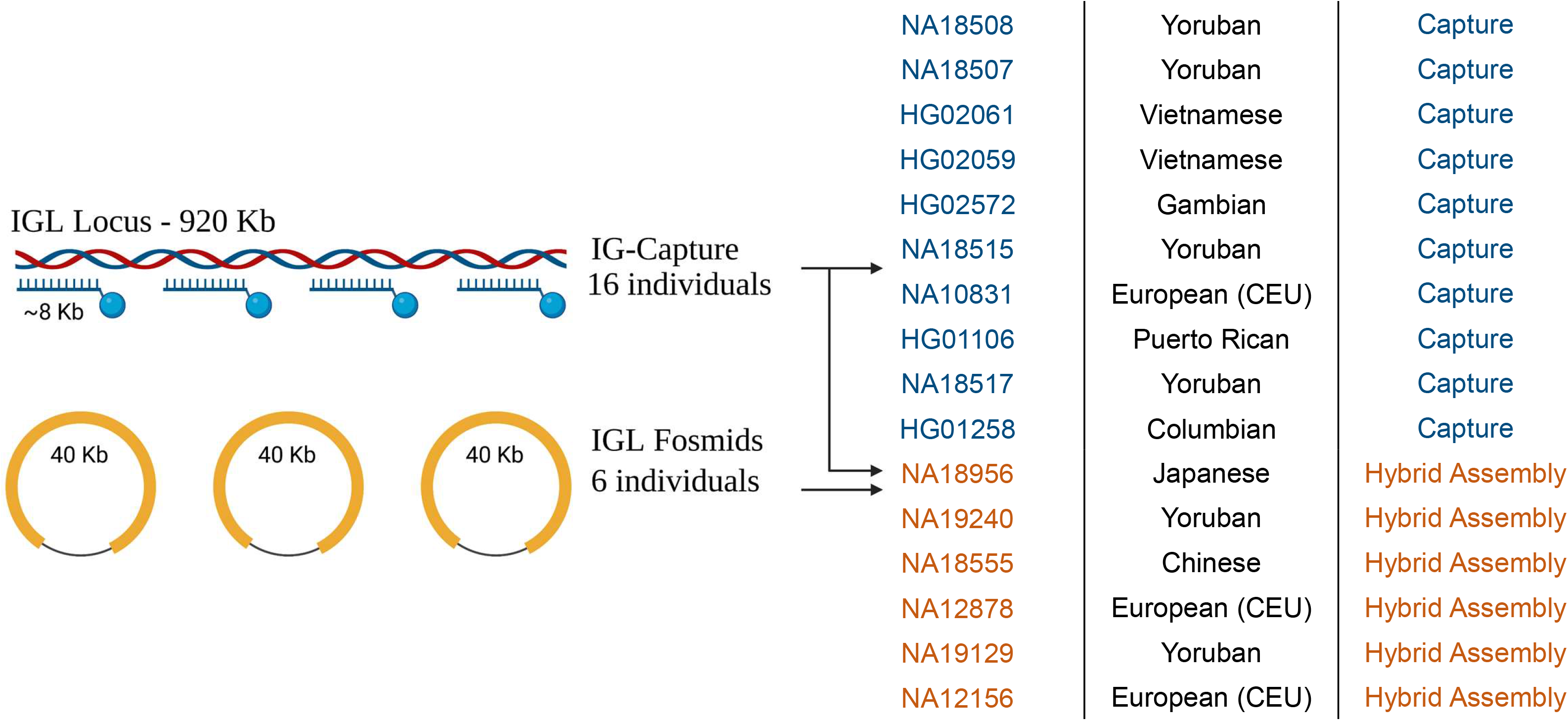

