## Supplemental Figures and Tables for "Characterization of the immunoglobulin lambda chain locus from diverse populations reveals extensive genetic variation"

#### Supplementary Tables and Figures

William S. Gibson, Oscar L. Rodriguez, Kaitlyn Shields, Catherine A. Silver, Matthew Emery, Gintaras Deikus, Robert Sebra, Evan Eichler, Ali Bashir, Melissa L. Smith*, Corey T. Watson*

### Supplementary Tables

| Sample | Haplotype | % GRCh38 Coverage | Assembly Coverage (bp) | No Coverage (bp) | IGL Size (bp) | Gaps Due to VDJ (bp) | Assembly Gaps (bp) |
|---|---|---|---|---|---|---|---|
| NA18956 | 1 | 100.00% | 920001 | 0 | 920001 | 0 | 0 |
| NA18956 | 2 | 100.00% | 920001 | 0 | 920001 | 0 | 0 |
| NA19240 | 1 | 100.00% | 920001 | 0 | 920001 | 0 | 0 |
| NA19240 | 2 | 100.00% | 920001 | 0 | 920001 | 0 | 0 |
| NA18555 | 1 | 100.00% | 920001 | 0 | 920001 | 0 | 0 |
| NA18555 | 2 | 100.00% | 920001 | 0 | 920001 | 0 | 0 |
| NA12878 | 1 | 94.70% | 871271 | 48730 | 920001 | 21192 | 0 |
| NA12878 | 2 | 20.02% | 184174 | 735827 | 920001 | 735827 | 0 |
| NA19129 | 1 | 99.11% | 911840 | 8161 | 920001 | 0 | 8161 |
| NA19129 | 2 | 99.53% | 915633 | 4368 | 920001 | 0 | 4368 |
| NA12156 | 1 | 100.00% | 920001 | 0 | 920001 | 0 | 0 |
| NA12156 | 2 | 98.28% | 904,147 | 15854 | 920001 | 0 | 15854 |

Table S1. Description of hybrid assembly coverage across the IGL locus, the size of assembly gaps due to VDJ artifacts, and the size of assembly gaps due to missing sequence. Percent GRCh38 coverage corresponds to the percentage of the GRCh38 reference that is overlapped by the resolved haplotypes.

| Sample | Average Coverage Depth | > 5x Coverage (bp) | No Coverage (bp) | Total Bases | Percent Coverag e | Haplotype gaps due to VDJ (bp) |
|---|---|---|---|---|---|---|
| NA18508 | 453 | 917345 | 501 | 920001 | 99.95% | 0 |
| NA18507 | 233 | 908674 | 1497 | 920001 | 99.84% | 0 |
| HG02061 | 150 | 914590 | 175 | 920001 | 99.98% | 108068 |
| HG02059 | 188 | 902666 | 1984 | 920001 | 99.78% | 0 |
| HG02572 | 143 | 902278 | 601 | 920001 | 99.93% | 0 |
| NA18515 | 141 | 909597 | 0 | 920001 | 100.00% | 0 |
| NA10831 | 142 | 907786 | 73 | 920001 | 99.99% | 203649 |
| NA18517 | 164 | 896823 | 9338 | 920001 | 98.97% | 0 |
| HG01258 | 111 | 891385 | 1730 | 920001 | 99.81% | 0 |
| HG01106 | 105 | 891293 | 2087 | 920001 | 99.77% | 0 |

| | **Average HiFi Depth** | | **Average Assembly Gaps (bp)** | **Avg Assembly % Coverage** |
|---|---|---|---|---|
| | 183.4 | | 1799 | 99.8% |

Table S2. Description of CCS read coverage across IGL locus. Note that VDJ gaps in HG02061 and NA10831 impact only a single haplotype.

| Sample | Minimum Block Size (bp) | Maximum Block Size (bp) | Average Block Size (bp) | Number of Phased Blocks | Percentage of IGL Locus Phased |
|---|---|---|---|---|---|
| NA18508 | 18683 | 196603 | 119027 | 8 | 93.93% |
| NA18507 | 25345 | 214336 | 118531 | 8 | 91.18% |
| HG02061 | 353 | 183431 | 41430 | 12 | 49.05% |
| HG02059 | 1848 | 244707 | 56290 | 9 | 66.87% |
| HG02572 | 20594 | 208682 | 68982 | 9 | 79.72% |
| NA18515 | 5396 | 206836 | 106707 | 9 | 89.33% |
| NA10831 | 2049 | 151719 | 48480 | 10 | 37.55% |
| NA18517 | 2023 | 198913 | 76700 | 10 | 93.00% |
| HG01258 | 3306 | 94095 | 36019 | 14 | 61.84% |
| HG01106 | 1699 | 200175 | 48513 | 12 | 78.10% |

Table S3. Description of phased assembly blocks across the IG regions for capture assemblies. IG regions are defined as regions unmasked in Fig 2.(A) and IGLC (chr22:22886736-22944092). Percentage of IGL Locus Phased refers to regions where both haplotypes are resolved and does not include large homozygous regions that cannot be split into haplotypes. Note that HG02061 and NA10831 have VDJ artifacts which render a portion of one haplotype unresolvable.

| Sample | HiFi Coverage | gene | Closest IMGT Allele | Percent Identity | Notes | AA differences | AA position |
|---|---|---|---|---|---|---|---|
| HG01106 | 15 | IGLV2-11 | IGLV2-11*01 | 99.65 | | 0 | |
| HG02572 | 33 | IGLV2-33 | IGLV2-33*01 | 99.65 | ORF | 0 | |
| NA18507, HG02572 | 93,48 | IGLV7-46 | IGLV7-46*01 | 99.65 | | 0 | |
| NA18507, NA18515 | 54,36 | IGLV5-37 | IGLV5-37*01 | 99.67 | | 0 | |
| NA18507 | 37 | IGLV2-14 | IGLV2-14*03 | 99.65 | | 0 | |
| NA18508 | 84 | IGLV1-47 | IGLV1-47*01 | 99.65 | | 0 | |
| NA18508 | 102 | IGLV3-21 | IGLV3-21*03 | 99.66 | | 0 | |
| NA18517 | 63 | IGLV7-43 | IGLV7-43*01 | 99.65 | | 0 | |
| HG02059, NA10831, HG01258 | 46,62,27 | IGLV5-37 | IGLV5-37*01 | 99.67 | | 1 AA, CDR1 | g103>a, G35>S (+ + -) |
| HG02061 | 12 | IGLV4-3 | IGLV4-3*01 | 99.32 | | 1 AA, CDR2 | c192>a, S64>R (- - -) |
| NA18508 | 85 | IGLV4-60 | IGLV4-60*03 | 99.66 | | 1 AA, CDR2 | g176>a, S59>N (- - -) |
| NA18508, NA18507, HG02572 | 96,38,36 | IGLV1-36 | IGLV1-36*01 | 99.65 | | 1 AA, CDR3 | c330>g, S110>R (- - -) |
| HG02059 | 43 | IGLV3-16 | IGLV3-16*01 | 99.65 | | 1 AA, CDR3 | g332>c, G111>A (- + -) |
| NA18507, NA18517 | 73,54 | IGLV10-54 | IGLV10-54*02 | 99.65 | | 1 AA, CDR3 | t320>g, L107>W (+ - -) |
| NA18515, NA18517 | 23,26 | IGLV2-33 | IGLV2-33*01 | 99.65 | ORF | 1 AA, FR1 | g37>a, G13>R (- - -) |
| NA18507, HG02059, NA18515, NA10831, HG01258, HG01106 | 36,42,32,29,21,17 | IGLV3-25 | IGLV3-25*03 | 99.64 | | 1 AA, FR1 | g74>c, G25>A (- + -) |
| HG01258 | 27 | IGLV3-22 | IGLV3-22*03 | 99.28 | | 1 AA, FR1 | t37>g, L13>V (+ - +) |
| NA18507 | 61 | IGLV2-18 | IGLV2-18*02 | 99.65 | | 1 AA, FR2 | a142>g, T48>A (- - -) |
| NA18507 | 92 | IGLV7-43 | IGLV7-43*01 | 99.65 | | 1 AA, FR2 | c155>t, A52>V (+ - +) |
| HG01106 | 22 | IGLV3-10 | IGLV3-10*01 | 99.64 | | 1 AA, FR2 | g115>c, A39>P (- - -) |
| NA18508, HG02572, NA18515 | 106,30,22 | IGLV3-16 | IGLV3-16*01 | 99.28 | | 1 AA, FR2 | t146>c, F49>A (+ - -) |
| HG02059 | 44 | IGLV1-50 | IGLV1-50*01 | 99.65 | ORF | 1 AA, FR3 | a197>g, N66>S (- - -) |
| NA18508, NA18507, NA18517 | 82,72,26 | IGLV1-50 | IGLV1-50*01 | 99.65 | ORF | 1 AA, FR3 | a224>g, Q75>R (+ - -) |
| HG02061, HG02059 | 27,72 | IGLV4-60 | IGLV4-60*03 | 99.66 | | 1 AA, FR3 | c259>t, R87>C (- - -) |
| HG02572 | 59 | IGLV10-54 | IGLV10-54*04 | 99.3 | | 1 AA, FR3 | g298>c, A100>P (- - -) |
| HG02061 | 39 | IGLV2-23 | IGLV2-23*02 | 99.31 | | 2 AA, CDR1, FR2 | g104>c, S35>T (+ - +) ; g140>a, G47>D (- - -) |
| NA18517 | 55 | IGLV4-3 | IGLV4-3*01 | 99.32 | | 2 AA, FR1, FR2 | c78>a, S26>R (- - -) ; c127>g, Q43>E (+ + -) |
| HG01258 | 12 | IGLV2-11 | IGLV2-11*01 | 99.3 | | 2 AA, FR2, FR2 | g132>c, Q44>H (- + -) ; t162>g, I54>M (+ + -) |
| NA18508, NA18515, NA18517 | 131,33,54 | IGLV2-23 | IGLV2-23*02 | 97.92 | | 3 AA, CDR1, CDR1, CDR2 | a103>g, S35>G (+ + -) ; c112>t, L38>Y (- - -) ; g168>t, E56>D (+ - +) |
| NA18508, HG02059, HG02572 | 139,61,40 | IGLV2-14 | IGLV2-14*01 | 99.31 | | 3 AA, CDR2, FR3, CDR3 | g168>t, E56>D (+ - +) ; t198>g, N66>K (+ - -) ; c337>t, L113>F (+ - -) |
| HG01258 | 19 | IGLV1-47 | IGLV1-47*01 | 99.65 | 1 bp insertion | frameshift (stop, stop , stop) | |
| HG02059 | 45 | IGLV3-22 | IGLV3-22*03 | 97.85 | 5 bp insertion | frameshift (stop, stop , stop) | |
| NA18508 | 135 | IGLV3-22 | IGLV3-22*03 | 98.57 | 5 bp insinsertion | frameshift (stop, stop , stop) | |

Table S4. Novel allele information. Regions with AA differences are based on IMGT annotations. AA position annotations were generated using IMGT V-Quest.

| Individual | IGLV5-39 Presence |
|---|---|
| HG02059 | 1 Allele |
| HG02061 | 2 Alleles |
| HG01258 | 2 Alleles |
| NA18515 | 1 Allele |
| NA18517 | 2 Alleles |
| NA10831 | 2 Alleles |
| HG01106 | 1 Allele |
| NA19240 | 2 Alleles |
| NA12156 | 2 Alleles |

Table S5. Individuals with 9.1 Kb IGLV5-39 insertions by number of haplotypes with the insertion allele.

| Individual | IGLV1-47 adjacent | IGLV8-61 distal |
|---|---|---|
| NA18956 | None | None |
| NA19240 | 1 Allele | None |
| NA18555 | 1 Allele | 1 Allele |
| NA12878 | None | None |
| NA19129 | None | None |
| NA12156 | 1 Allele | 1 Allele |
| NA18508 | None | None |
| NA18507 | None | 1 Allele |
| HG02061 | 1 Allele | 1 Allele |
| HG02059 | 2 Alleles | None |
| HG02572 | None | None |
| NA18515 | None | None |
| NA10831 | 1 Allele | None |
| NA18517 | None | 1 Allele |
| HG01258 | 1 Allele | 1 Allele |
| HG01106 | 1 Allele | 1 Allele |

Table S6. Individuals with LINE insertions by location and number of haplotypes with the insertion allele.

| Location (GRCh38) | Adjacent Genes | Size (bp) | Characteristics |
|---|---|---|---|
| chr22:22,361,235-22,361,274 | IGLV1-47 | ~6,000 | LINE insertion |
| chr22:22,131,956-22,131,995 | IGLV4-60, IGLV8-61 | ~6,000 | LINE insertion |
| chr22:22,875,333-22,875,456 | IGLV3-1, IGLV3-2 | 250 | 35 bp tandem repeat |
| chr22:22,585,771-22,586,112 | IGLV2-33, IGLV2-34 | 230 | Short tandem repeat |
| chr22:22,183,041-22,183,080 | IGLV(V)-58 | 175 | 36 bp tandem repeats |
| chr22:22,179,716-22,179,755 | IGLV(IV)-59 | 320 | 19-21 bp poly-A repeat |

Table S7. Table of structural variations in the IGLV region which do not contain genes.

| Sample | Homozygous Alt | Heterozygous | Total SNVs | IG-only Homozygous Alt | IG-only Heterozygous | Total SNVs IG-only | Percentage of Homozygous Alt SNVs in IG-only regions | Population |
|---|---|---|---|---|---|---|---|---|
| NA18956 | 1506 | 948 | 2454 | 987 | 535 | 1522 | 64.85% | JPN |
| NA19240 | 1142 | 1772 | 2914 | 664 | 994 | 1658 | 40.05% | YRI |
| NA18555 | 1138 | 1785 | 2923 | 698 | 1102 | 1800 | 38.78% | CBG |
| NA12878 | 1320 | 299 | 1619 | 768 | 193 | 961 | 79.92% | CEU |
| NA19129 | 880 | 1995 | 2875 | 457 | 1168 | 1625 | 28.12% | YRI |
| NA12156 | 689 | 1333 | 2022 | 412 | 659 | 1071 | 38.47% | CEU |
| NA18508 | 1029 | 1976 | 3005 | 518 | 1262 | 1780 | 29.10% | YRI |
| NA18507 | 970 | 1973 | 2943 | 638 | 1085 | 1723 | 37.03% | YRI |
| HG02061 | 857 | 1272 | 2129 | 646 | 686 | 1332 | 48.50% | KHV |
| HG02059 | 1025 | 1699 | 2724 | 707 | 839 | 1546 | 45.73% | KHV |
| HG02572 | 588 | 1325 | 1913 | 334 | 871 | 1205 | 27.72% | GWD |
| NA18515 | 822 | 1592 | 2414 | 414 | 1017 | 1431 | 28.93% | YRI |
| NA10831 | 556 | 643 | 1199 | 466 | 243 | 709 | 65.73% | CEU |
| NA18517 | 611 | 1767 | 2378 | 295 | 1180 | 1475 | 20.00% | YRI |
| HG01258 | 404 | 947 | 1351 | 248 | 620 | 868 | 28.57% | CLM |
| HG01106 | 409 | 993 | 1402 | 203 | 710 | 913 | 22.23% | PUR |

Table S8. Total SNV counts and SNV counts split by heterozygous or homozygous alternate for the IGLV region (chr22:22024092-22886736) and the IG-only region (see Fig. 2A).

A. Across IGL locus (chr22:22,024,092-22,944,092)

| Unique SNVs | Multiallelic SNVs | SNVs not found in dbSNP "All" | SNVs not found in dbSNP "Common" |
|---|---|---|---|
| 7599 | 39 | 381 | 2672 |

B. Across IG-only regions (see Figure 2A) in IGLV (chr22:22024092-22886736)

| Unique SNVs | Multiallelic SNVs | SNVs not found in dbSNP "All" | SNVs not found in dbSNP "Common" |
|---|---|---|---|
| 4298 | 23 | 124 | 939 |

Table S9. Table of unique SNVs across 13 individuals, V(D)J artifact individuals are excluded. (A) Across the entire IGL locus, unique and multiallelic SNVs, SNVs not found in dbSNP “All” or “Common” as of 2022/03/23. (B) Across only IG-regions in the IGLV locus, unique and multiallelic SNVs, SNVs not found in dbSNP “All” or “Common” as of 2022/03/23.

| IGLJ-C Cassette | Minimum Similarity | Maximum Similarity |
|---|---|---|
| IGLJ-C1 | 99.183% | 99.981% |
| IGLJ-C2 | 98.722% | 100% |
| IGLJ-C3 | 98.025% | 100% |
| IGLJ-C4 | 99.621% | 99.976% |
| IGLJ-C5 | 99.67% | 100% |
| IGLJ-C6 | 99.334% | 100% |
| IGLJ-C7 | 99.203% | 100% |

Table S10. Sequence of each IGLJ-C cassette group for all haplotypes was compared. The maximum similarity is the percent similarity of the closest matching haplotypes, the minimum similarity is the percent similarity of the least matching haplotypes.

### Supplementary Figures

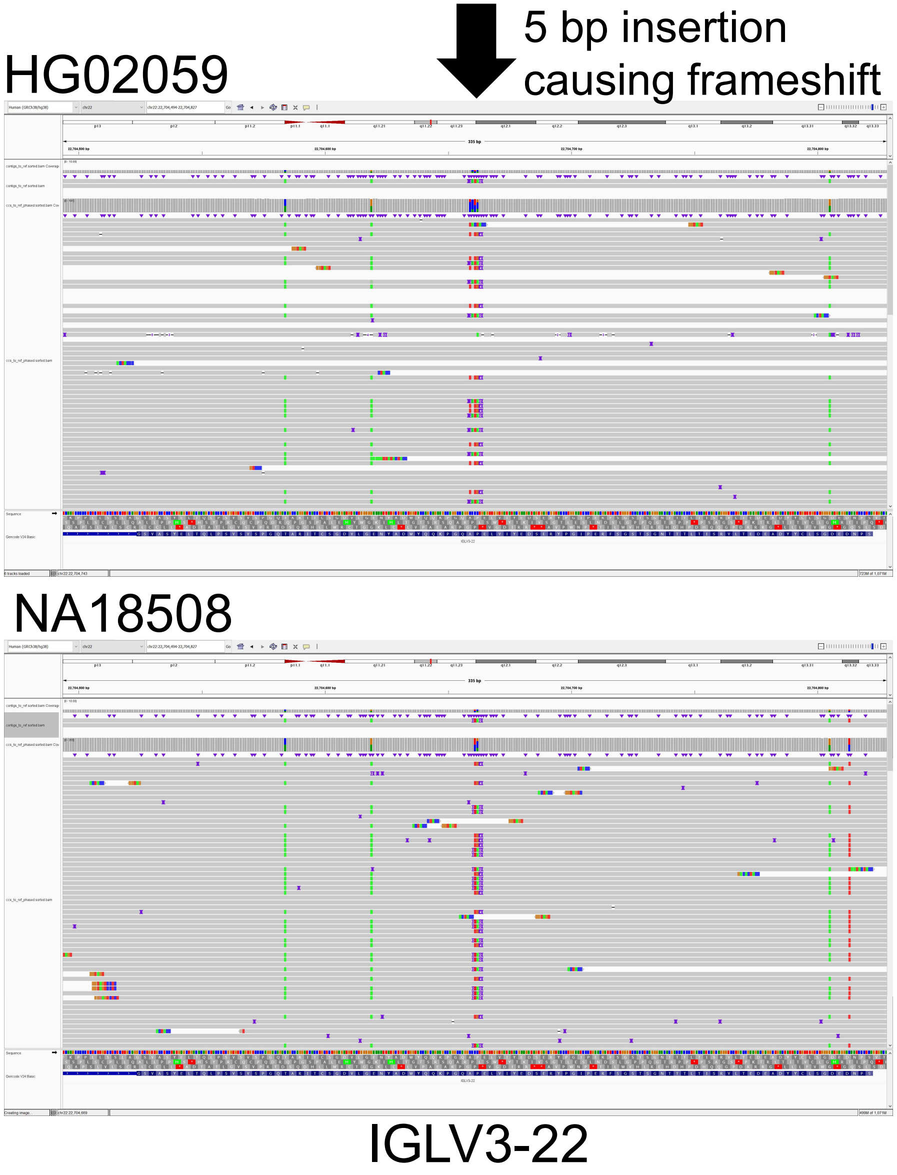

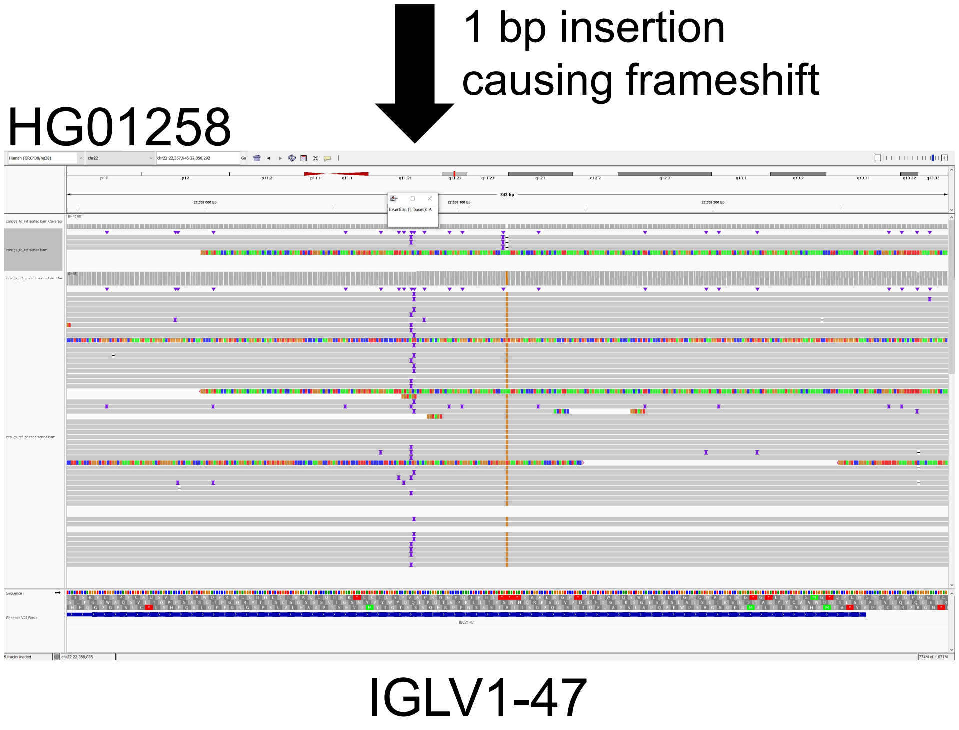

Figure S1. Novel alleles with frameshifts resulting in nonfunctional genes confirmed with HiFi reads showing nucleotide insertions.

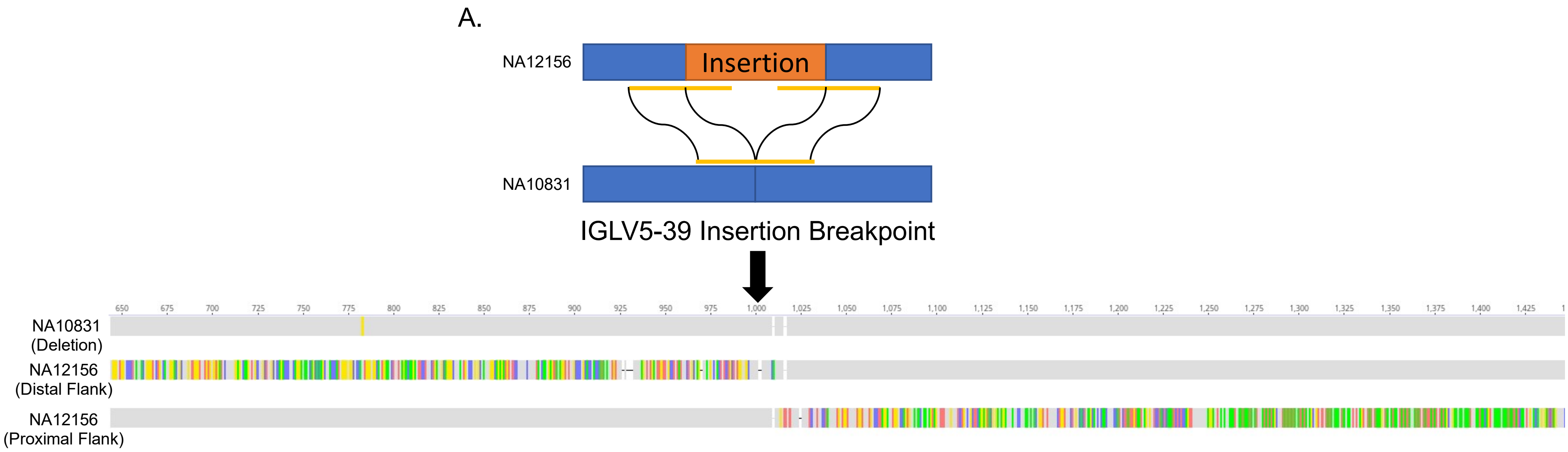

Figure S2. A.1 Kb of sequence flanking the IGLV5-39 insertion breakpoints of NA12156 were compared to 2 Kb of the same region from NA10831 which has IGLV5-39 deleted. The 1 Kb proximal and distal flanks of the IGLV5-39 insertion from NA12156 are highly homologous to NA10831 while the 1 Kb sequence inside the IGLV5-39 insertion differs significantly.

### IGLV5-39 SV AT-rich region

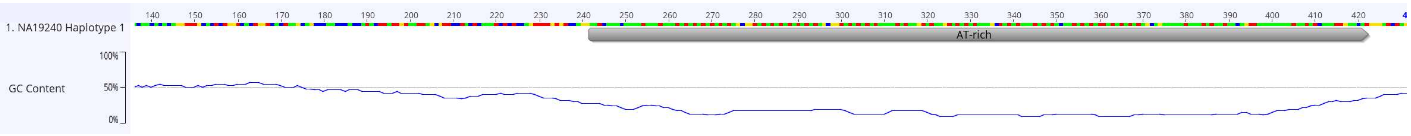

| | Freq | % |
|---|---|---|
| A: | 58 | 32.0% |
| C: | 10 | 5.5% |
| G: | 13 | 7.2% |
| T: | 100 | 55.2% |
| GC: | 23 | 12.7% |
| All: | 181 | 100.0% |

Figure S3. AT-rich region adjacent to breakpoint in IGLV5-39 SV. The annotated AT-rich region has 87.3% A/T content.

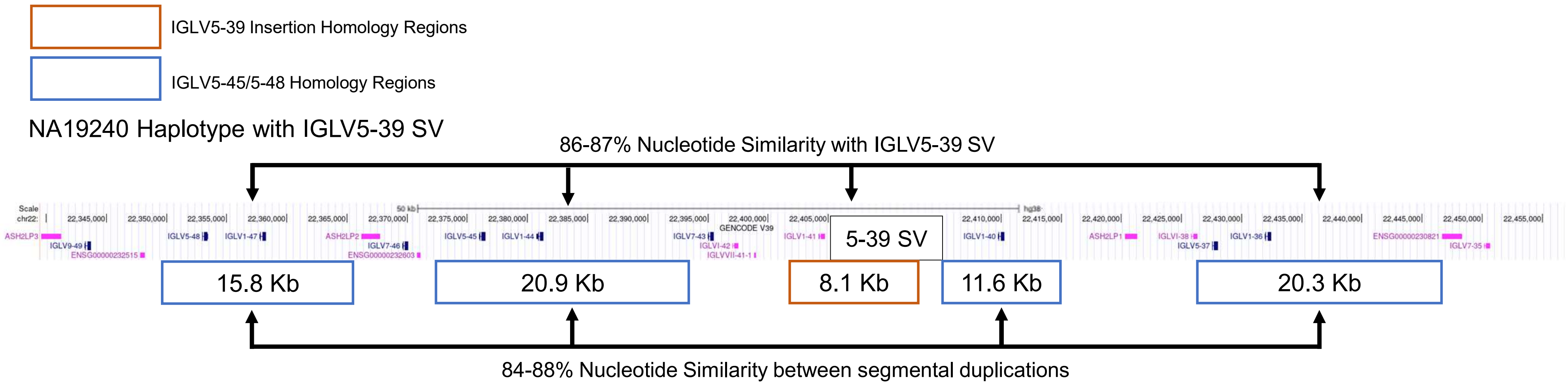

Figure S4. The NA19240 haplotype with the IGLV5-39 SV is shown. The orange box is the IGLV5-39 SV segmental duplication region. The blue boxes are regions of segmental duplication sequence surrounding the IGLV5-37, IGLV5-45, and IGLV5-48 genes.

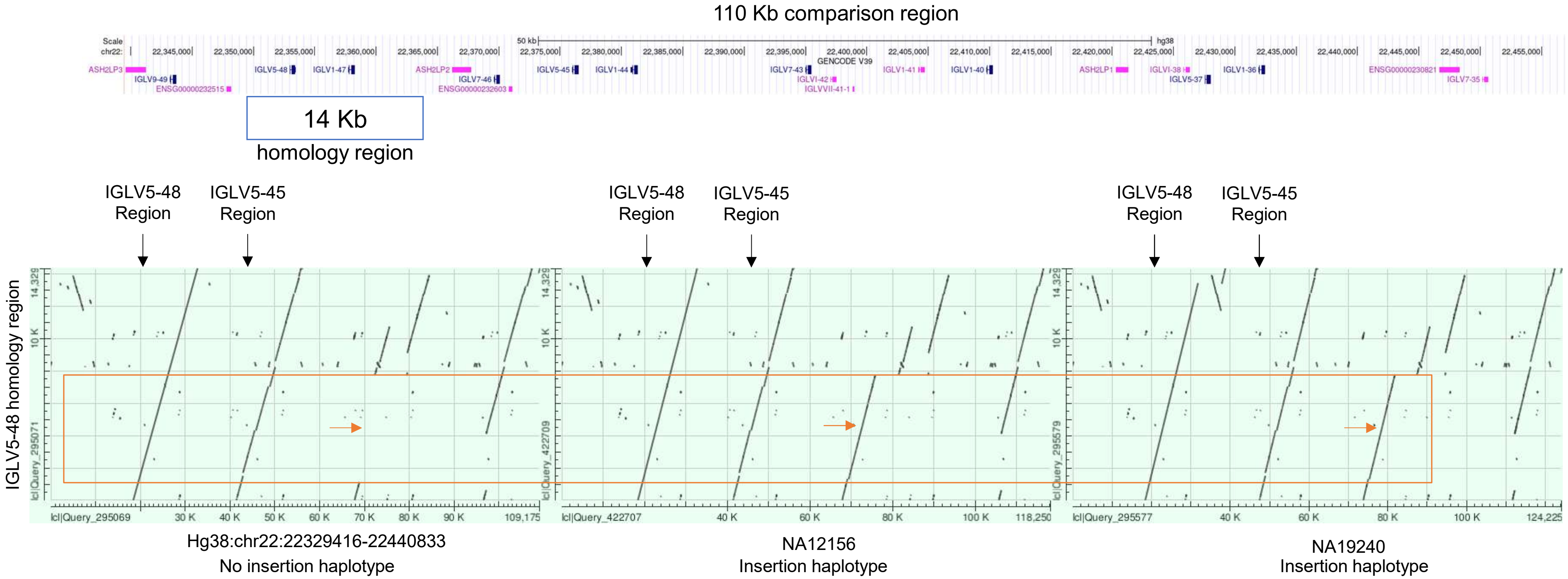

Figure S5. IGLV5-48 homology region comparison to Non-insertion and Insertion Haplotypes. Each dotplot has a 14 Kb homologous region surrounding IGLV5-48 on the Y-axis. The first dotplot has the Hg38 reference (lacking the IGLV5-39 insertion) on the X-axis, the other two dotplots are of the same region on the X-axis but both contain the IGLV5-39 reference (NA12156 and NA19240). The orange box highlights the region of homology between the IGLV5-48 region and the IGLV5-39 insertion. The orange arrows are pointing to the region where the insertion does/does not reside. The black arrows are denoting the IGLV5-45/5-48 homologous regions.

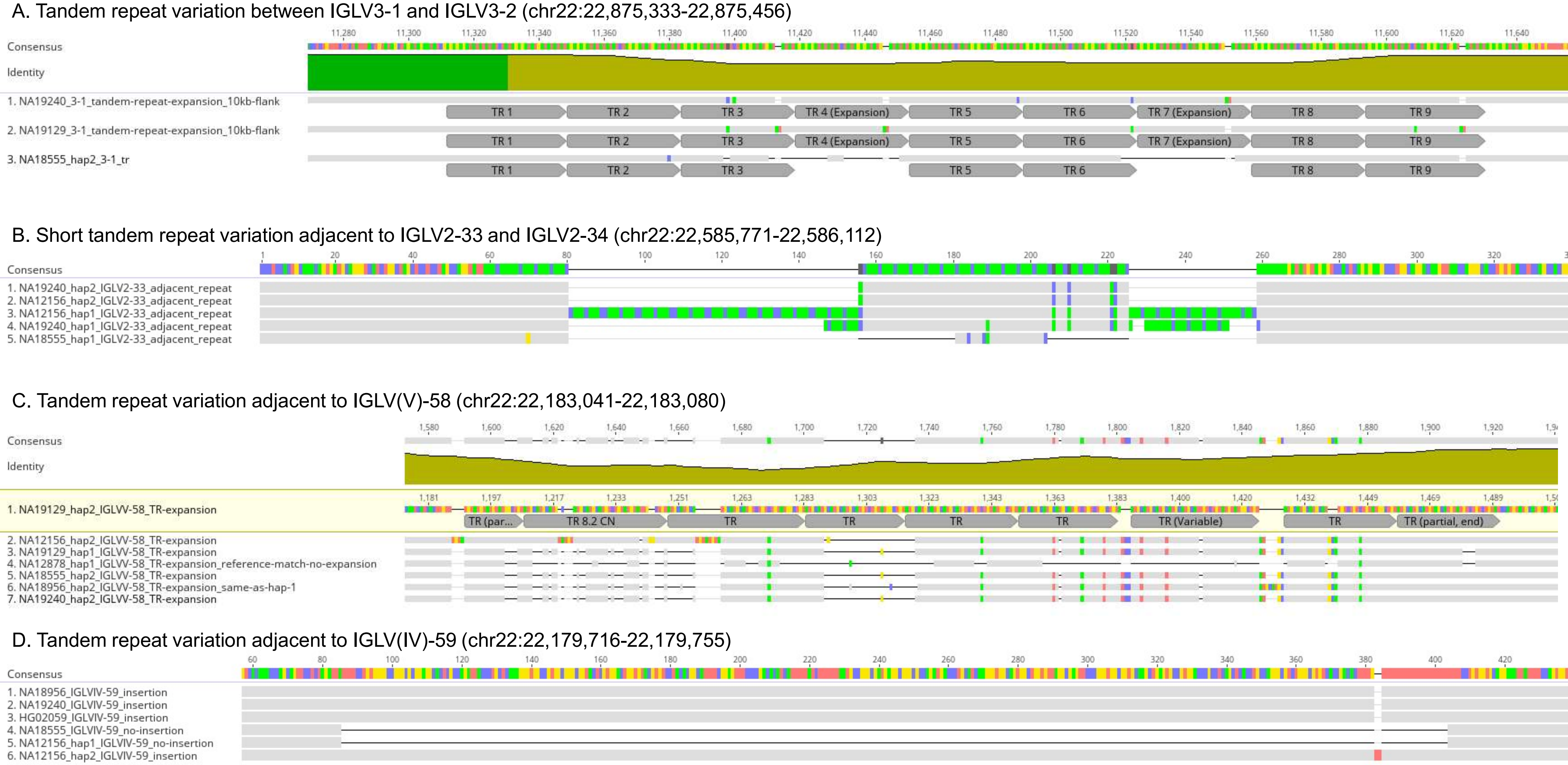

Figure S6. (A) Variation in a ~35 bp tandem repeat between IGLV3-1 and IGLV3-2 (chr22:22,875,333-22,875,456). (B) Variation in a short tandem repeat (CTTT) between IGLV2-33 and IGLV2-34 (chr22:22,585,771-22,586,112). (C) Variation in a ~36 bp tandem repeat adjacent to IGLV(V)-58 (chr22:22,183,041-22,183,080). (D) Insertion of 316-318 bp adjacent to IGLV(IV)-59 (chr22:22,179,716-22,179,755)

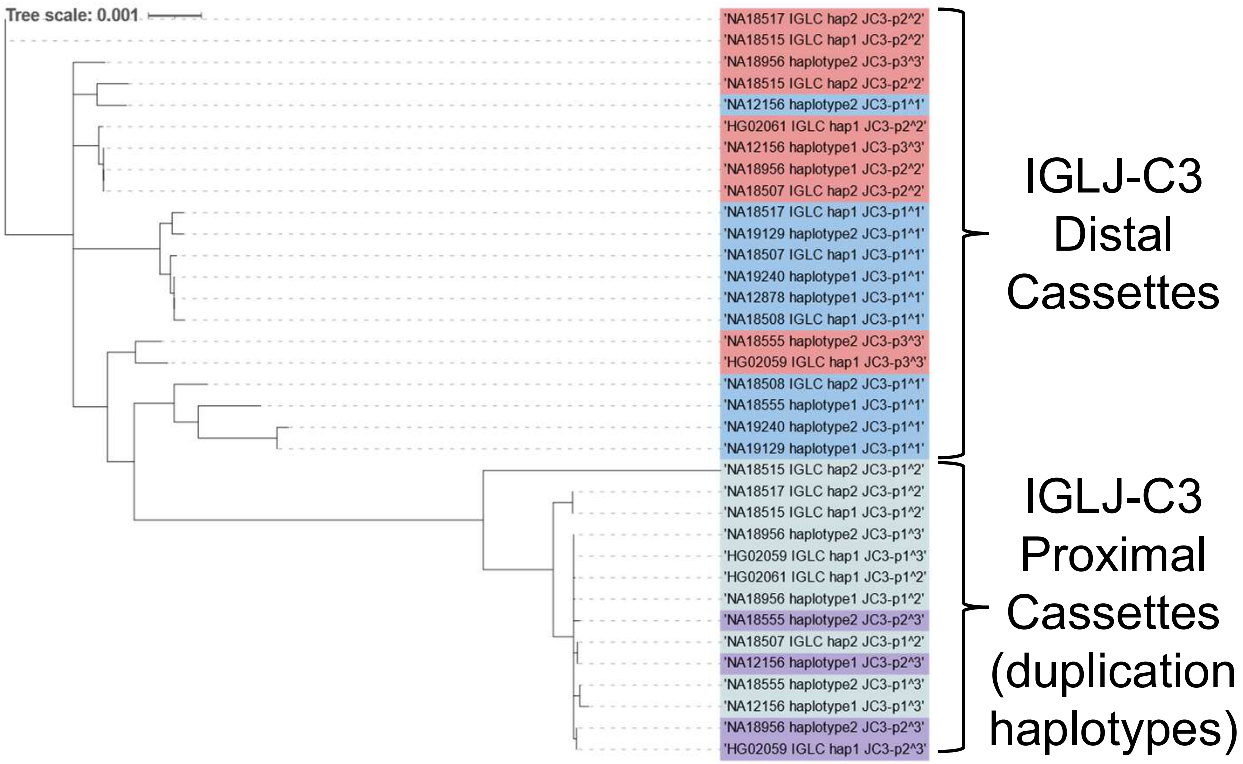

Figure S7. Phylogeny of the entire IGLJ-C3 cassettes. Colors indicate if the cassette is part of a duplication haplotype and where the cassette exists. (Blue = No duplication haplotype ; Red = Duplication haplotype bordering IGLJ-C4 ; Grey = Duplication haplotype bordering IGLJ-C2 ; Purple = Duplication haplotype in between two IGLJ-C3 cassettes). Proximal cassettes border IGLJ-C2 and a second IGLJ-C3 cassette. Distal cassettes border IGLJ-C4 and can be from duplication or non-duplication haplotypes.

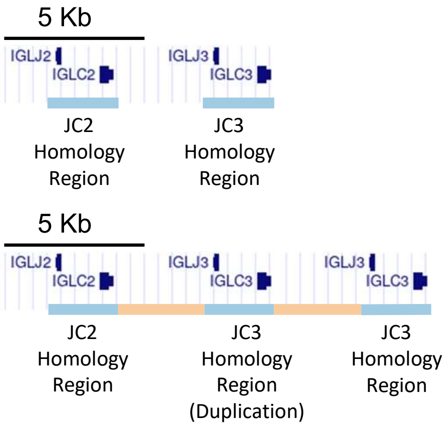

Figure S8. The blue bars in the first panel show highly homologous segments (>99%) between the IGLJ-C2 and IGLJ-C3 cassettes. These homology regions are compared in Fig. S9. The second panel shows the 2 KB homologous regions (blue) with additional homologous regions (orange) in the context of IGLJ-C duplication. The orange regions also have >97% identity with each other.

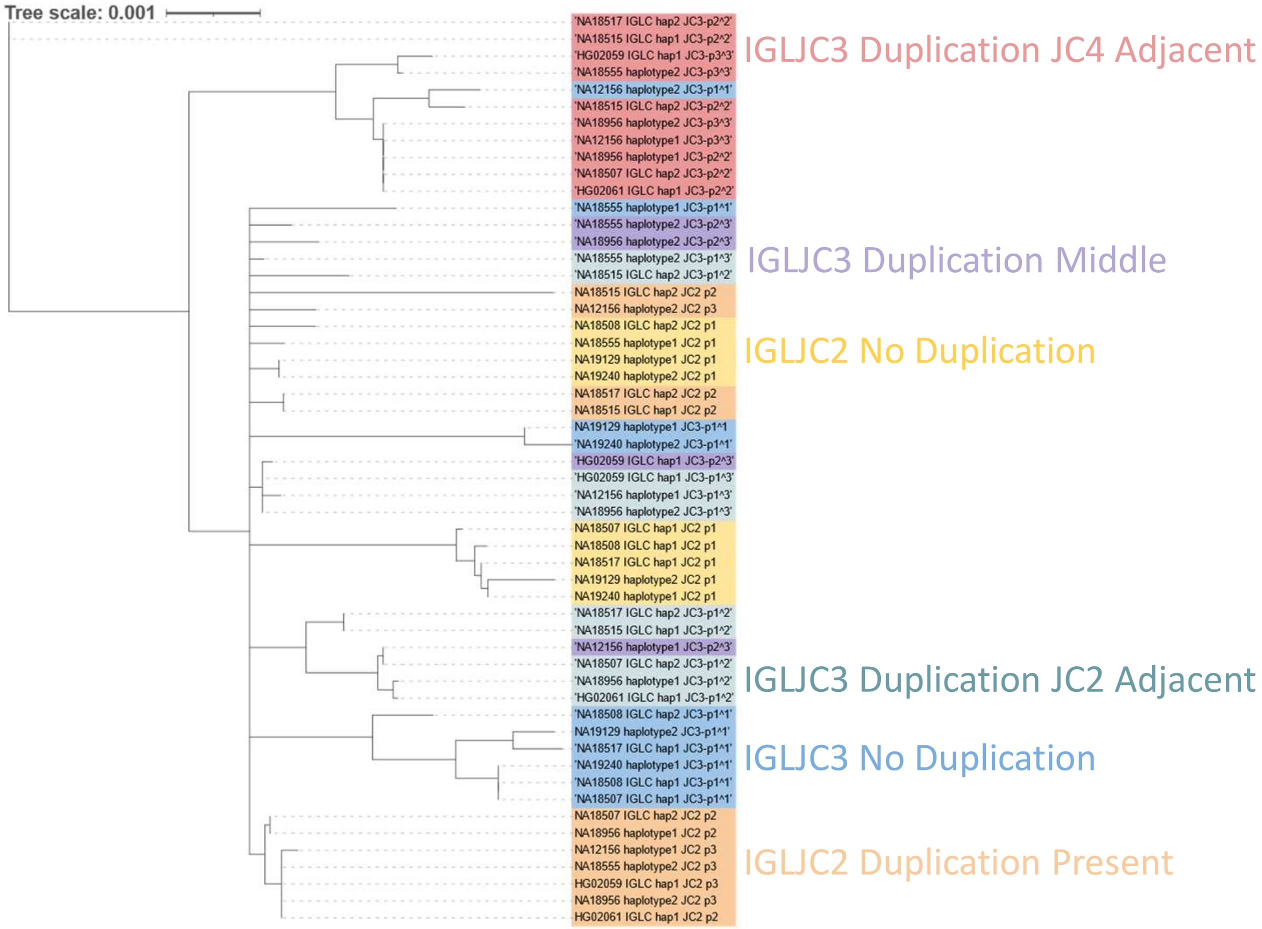

Figure S9. Phylogenetic analyses of 2 KB homology regions for each IGLJ-C2 and IGLJ-C3 haplotype. Colors indicate if the homology region is part of a duplication haplotype and if so where in the haplotype it exists relative to the other cassettes. 'Duplication JC2 Adjacent' is the proximal-most IGLJ-C3 duplication, 'Duplication JC4 Adjacent' is the distal-most IGLJ-C3 duplication, 'Duplication Middle' is the middle IGLJ-C3 duplication in the case of three IGLJ-C3 copies.
